# *Ambecovirus*, a novel *Betacoronavirus* subgenus circulating in neotropical bats sheds new light on bat-borne coronaviruses evolution

**DOI:** 10.1101/2025.07.20.665586

**Authors:** Gabriel da Luz Wallau, Eder Barbier, Lais Ceschini Machado, Alexandre Freitas da Silva, Yago Jose Mariz Dias, Alexandru Tomazatos, Balázs Horváth, Roberto D. Lins, Enrico Bernard, Dániel Cadar

## Abstract

Understanding the viral diversity harboured by wildlife is essential for effective prediction and prevention of future zoonotic outbreaks. Bats, in particular, are recognized as natural reservoirs for several zoonotic viral pathogens of high impact on public health, including coronaviruses responsible for SARS, the rabies virus, Marburg, Ebola, Nipah and Hendra viruses. However, the large extent of bat viruses remains unexplored, especially in highly biodiverse regions of the Neotropics such as Brazil. We used a meta-transcriptomic to characterize new virus genomes found in blood, oral and anal samples collected from cave- and non-cave bats from Northeast Brazil. From a total of 19 coronavirus-positive bats, we have assembled two complete genomes of a new *Betacoronavirus* subgenus, named *Ambecovirus* (American betacoronavirus). The subgenus herein described is phylogenetically placed between the *Sarbeco*-/*Hibeco*-/*Nebeco*virus and the *Merbeco*-/*Embecovirus* clades, being basal to the former. While the conserved S2 region of the spike protein retained hallmark domains, including HR1 and HR2, the S1/S2 cleavage site and the furin cleavage site, the S1 region consistently displayed only the N-terminal domain. The receptor-binding domain could not be identified due to high dissimilarity relative to known congeneres. The detection of *Ambercovirus* in sympatric *Pteronotus gymnonotus* and *Carollia perspicillata* bats suggests interspecies transmission. Longitudinal sampling confirmed persistent *Ambecovirus* infection in *P. gymnonotus* over multiple years and virus dispersion at a minimum distance of 270 km between caves. The present study confirms that viral diversity in neotropical hosts remains largely unknown not just in Brazil but, likely, in the other countries of the region, supporting the need for a systematic approach to virome exploration and analysis followed by *in vitro* experimentation to assess zoonotic potential.

## Introduction

Viruses are obligate parasites and represent some of the most widespread and biologically diverse entities around the planet (1,2). Due to this parasitic lifestyle, many viruses cause diseases in their hosts, but as our knowledge advances on the virosphere (viral communities), evidence points to a consensus that the majority of viruses are non-pathogenic with a small fraction with clear pathogenic profile (3–5). Still, such evidence is based on a very limited - but expanding - view about the known virosphere (3,6). Regarding pathogenic viruses, a growing body of evidence suggests that several natural host reservoirs are tolerant to infections and experience little or no disease likely due to the long-term host-virus coevolution, and viral tolerance profiles evolved along millions of years of evolution (7). Bats (Order Chiroptera), the only mammals capable of sustained flight, are special examples of reservoirs of a number of known pathogenic viruses and, yet, appear mostly lightly or even unaffected by them (8). This group of mammals have gone through multiple adaptations to perform active flight, including evolving differential immunological responses, which are linked to their virus tolerance profile (9,10).

Bats are the second largest order of mammals (~22% of known species) and have a worldwide distribution (11). Aside from their ecological roles as keystone-species (e.g. in plant pollination and seed dispersal), they are recognized as natural reservoir hosts of several viruses of high zoonotic importance that belong to different viral families such as *Coronaviridae*, *Paramyxoviridae*, *Rhabdoviridae* and *Filoviridae* (12). Our current understanding of their role in maintaining the circulation of several zoonotic virus species relies mainly on longitudinal surveillance efforts focused on rabies, Hendra and Nipah viruses in the Americas, Europe, Asia and Oceania (13,14). More recently, it also became evident that bats are the natural reservoirs of coronaviruses, including close relatives of human pathogens like SARS-CoV, SARS-CoV-2 and MERS-CoV betacoronaviruses, as well as of other *alphacoronaviruses*, such as HCoV-229E and HCoV-NL63 associated with seasonal, mild respiratory disease in humans (15–18). Most of the evidence on bats as natural reservoirs of these viruses comes primarily from low-throughput molecular methods that can capture a limited amount of the viral genetic diversity (14,19). More recently, unbiased high-throughput sequencing (metagenomics and meta-transcriptomics) has reshaped our understanding of viromes hosted by bats and by other host species, allowing hypothesis-free explorations without *a priori* knowledge of existing genetic diversity (5,20). These studies have corroborated many findings about main reservoir groups, but also uncovered new host groups and a hidden diversity of viral lineages (6).

Brazil is ranked first among the 17 megadiverse countries (21), covering six terrestrial biomes and the highest number of endemic species. With yet-uncharacterized viromes and microbiomes, plus an increasing anthropogenic pressure represented by deforestation and human encroachment in once pristine ecosystems, Brazil has been ranked as one of the most important potential hotspots for the emergence of new viral pathogens (22–25). With 186 known species in nine families, Brazil accounts for 12.5% of the world’s bat fauna (1.487 species) (26). Despite such bat species richness and diversity, nearly half of all bat species in Brazil have not been yet investigated as virus hosts (19). Moreover, most of the virus detection research performed so far in the country relied mainly on low throughput and targeted methodologies. Thus, the results obtained so far have been biased by preferential sampling of known reservoirs of rabies virus (19). Additionally, few longitudinal studies were performed using varied sampling strategies across different biomes in Brazil. Coronaviruses, for example, have been detected in neotropical bats from Mexico, Brazil, Argentina, Peru, Trinidad and Tobago and other countries, primarily through PCR and Sanger sequencing (27–37,37,38). These findings revealed a predominance of alpha-over betacoronaviruses infecting neotropical bats (39,40). So far, only one genomic study has been performed in Brazil reporting two betacoronaviruses, the first a MERS-CoV-related virus, while the second showed no clear phylogenetic clustering with known *Orthocoronavirinae* genera (41). These findings underscore the substantial gaps in our understanding of the bat virome and its zoonotic potential in Brazilian ecosystems as well as on coronavirus evolution.

Here we employed meta-transcriptomic untargeted high throughput sequencing for virus discovery in cave- and non-cave bats from Northeastern Brazil, particularly from the Caatinga drylands and from the Atlantic Forest biomes. We recovered complete *Betacoronavirus* genomes and compared their similarity to known coronaviruses using a comparative analysis to characterise the architecture of genomes, spike protein domains and revisit the deep evolutionary roots of the *Betacoronavirus* genus.

## Material and Methods

### Bat sampling

Sampling was performed from May 2019 until January 2023 across 12 sites in Northeastern Brazil (**Figure 1**). The initial collection, associated with a natural history and ecology survey, involved whole blood samples (30–50 µL) mixed with 100 µL of RNAlatter (Thermo Fisher Scientific). From 2020 onwards, oral and anal swabs were collected and directly enclosed in Eppendorf tubes and placed in liquid nitrogen containers without any buffer solution. Swabs were then transferred to −80°C freezers after nitrogen liquid containers reached stationary laboratories and remained there until sample processing.

**Figure 1.**
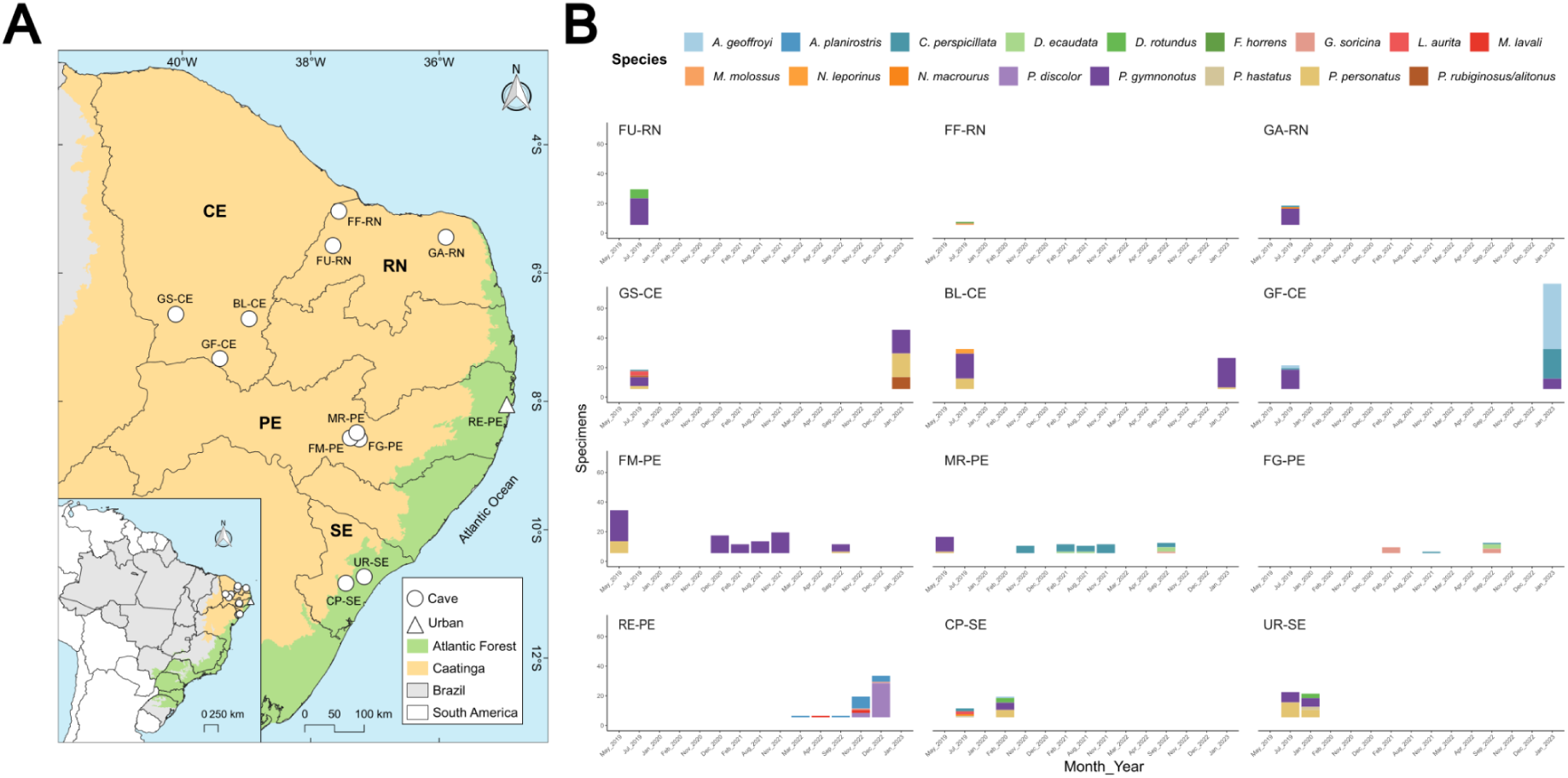
Bat sampling performed during the study period. **A** - Sampling sites including 11 caves (circles) and 1 urban (triangle) site in the Northeast part of Brazil. Colors denote two of the six Brazilian biomes; orange (Caatinga) and green (Atlantic forest). **B** - Number of specimens collected per sampling site during this study. The first two letters of the site acronym stands for the name of the cave (GS - Gruta do Sobradinho, BL - Boqueirão de Lavras II, GF - Gruta do Farias, FF - Furna Feia, FU - Furna do Urubu, GA - Gruta do Arnoud, MR - Meu Rei, FM - Furna do Morcego, FG - Furna do Gato, UR - Urubu, CP - Casa de Pedra) while the following two letters for stand for the state where the cave was located (CE - Ceará, RN - Rio Grande do Norte, PE - Pernambuco and SE - Sergipe).

Bats were captured using harp traps placed at cave entrances at 5:30 p.m. for one hour. In urban areas, bats were captured using four mist nets (12 m x 2.5 m each) set for a 4-hour period starting at dusk. Trapped/netted individuals were carefully removed, weighted and identified at the species level following specific literature (42–45). The bat’s age was estimated based on the metacarpal epiphyseal cartilages (46), with individuals classified as adults (closed epiphysis) or subadults (open epiphysis). The reproductive stage of bats was assessed based on the presence of secondary sexual characteristics. Female individuals were classified into three categories: pregnant, lactating, or inactive (non-reproductive). Males were considered reproductively active when they exhibited visibly enlarged testes. All bat handling was conducted by trained personnel with protection equipment, including NFP2 masks, face shield, gloves and lab coat. Fieldwork permit was issued by the Instituto Chico Mendes de Conservação da Biodiversidade–ICMBio through SISBIO authorizations (68992-1, 68992-2, 68992-3, 77600-1, 83959-1, and 83959-2) and approval by the Ethics Committee on Animal Care (CEUA) of the Universidade Federal de Pernambuco (Process numbers 114/2019 and 092/2021). Moreover, the study was also registered in Brazil’s Sistema Nacional de Gestão do Patrimônio Genético e do Conhecimento Tradicional Associado (SisGen) under the number A35D254.

### Sampling processing and metagenomic sequencing

We applied a virus-optimized metagenomic sequencing protocol adapted from established approaches for unbiased viral detection (46) to characterize the virome of the cave- and non-cave bats. Specimens were pooled by species, location, and collection date (1–4 individuals per pool). To reduce host and microbial nucleic acid background, samples underwent a viral particle enrichment protocol. Blood samples were diluted in sterile PBS, and swab samples were eluted in PBS, then incubated with Proteinase K at 50 °C for 30 minutes. The supernatant was subsequently filtered through a sterile 0.45 µm membrane (Millipore) to eliminate cellular debris and larger contaminants. Filtered aliquots (250–400 µL) were treated with a cocktail of nucleases, including Turbo DNase (Ambion), Baseline-ZERO DNase (Epicenter), Benzonase (Novagen), and RNAse One (Promega), to selectively degrade unprotected DNA and RNA, enriching for viral particles. Total nucleic acids were extracted using the MagMAX Viral RNA Isolation Kit (Life Technologies) for blood and the QIAamp Viral RNA Mini Kit (Qiagen) for swab samples, according to manufacturer protocols. Each sequencing batch included a negative water control to monitor for reagent contamination. Reverse transcription and cDNA synthesis were performed using an octamer random primer and the amplified DNA was used for library preparation with the QIAseq FX DNA Library Kit (Qiagen), incorporating dual-index barcoding to enable multiplex sequencing. Library concentrations were assessed using Qubit fluorometry and the Agilent Bioanalyzer. Sequencing was performed on the Illumina NextSeq 2000 platform (2×100 cycles).

### Bioinformatic analysis

#### Assembly and viral contigs binning

Raw sequences were filtered for quality (Phred score ≥20) and length (minimum 50 bases) and trimmed using Trimmomatic 0.39 to remove polyclonal and low-quality reads (47). To reduce artefactual read duplication introduced during random RT-PCR, deduplication was performed using the ‘dedupe.sh’ tool from the BBTools suite (v39.01) (48). Ribosomal RNA was removed by mapping clean, deduplicated reads to the SILVA database v138.1 (49). High quality, filtered sequences were then *de novo* assembled with Megahit v1.2.9 (50) with a minimum length limit of 400 bases. Contigs were compared against the NCBI non-redundant (nr) database (2024.03.16.) using DIAMOND v2.1.9 blastx with E-value cutoff of 0.001 for high sensitivity and to reduce false positives (51). To retrieve standardized assembly genomic metrics and consensus genomes, a complementary reference-based assembly approach was employed. This involved passing the newly complete and annotated genomes recovered from the *de novo* assembly strategy and the raw sequencing reads to ViralFlow v1.0 (52). CoverM tool (https://github.com/wwood/CoverM) was also used to obtain genomic metrics (53).

#### Contigs extension and annotation

The contigs were visually validated using the Geneious Prime 2024 mapping algorithm and extended through several iterations where possible. ORFs were predicted with Prodigal v2.6.3 (54) and MEGAN6 (55), then manually corrected as needed. Special attention was given to potential virus-specific coding strategies (e.g., ribosomal shifting, overlapping reading frames, transcriptional slippage, leaky scanning, alternative splicing) by using the closest available reference sequence in CLC Genomics Workbench 24 (Qiagen). All genomic sequences generated during this study have been submitted to GenBank and accession numbers will be updated here as soon as they become available. The associated raw sequencing datasets are publicly accessible via the Sequence Read Archive (SRA), linked to Bioproject ID PRJNA1291827.

#### Homologous sequences recovery

To investigate the evolutionary relationships of the identified coronaviruses, we performed homologous sequence searches using BLASTn and BLASTp at NCBI (last accessed June 2025). To place these new coronaviruses within the *Orthocoronavirinae* phylogenetic tree we used the amino acid sequences of the hallmark genes such as RNA dependent RNA polymerase (RdRp) for homologous searches. This included the amino acid sequence of reference viral genomes from the *Orthocoronavirinae* subfamily classified by the International Committee on Taxonomy of Viruses (https://ictv.global/). Phylogenetic reconstruction was based on RdRp sequences, since it reflects vertical ancestry and is less prone to recombination than other genomic regions (56). Additionally, to explore conserved domains in rapidly evolving proteins, we conducted amino acid homology searches of spike glycoproteins, which mediate host cell receptor binding. On the other hand, to study more recent evolutionary relationships we also performed blastn searches at the NCBI Virus (https://www.ncbi.nlm.nih.gov/labs/virus/vssi/#/) and the ZOVER databases (https://www.mgc.ac.cn/cgi-bin/ZOVER/main.cgi).

#### Multiple sequence alignment and phylogenetic reconstruction

Recovered and generated sequences were aligned using MAFFT v7 (57) and inspected using Aliview (58) both at the amino acid and nucleotide level. Phylogenetic reconstruction was performed using IQTREE v2 (59) after model selection with ModelFinder (60) implemented in IQTREE. Branch support was accessed using aLRT and ultrafast bootstrap (UFboot) with 1000 replicates. Beast v1.10 (61) was used to perform a Bayesian phylogenetic reconstruction for the *Orthocoronavirinae* phylogenetic tree to reassess specific branch support.

#### In silico protein characterization

In order to obtain more detailed insights about the betacoronavirus genome recovered we sought to characterize the conserved domains of the spike protein through searches at the Conserved Domain Database (CDD search - https://www.ncbi.nlm.nih.gov/Structure/cdd/wrpsb.cgi). To further explore and attempt to align the spike protein of the viruses sequenced in this study with other coronavirus spikes we also used the COBALT software for constraint-based alignment tool (62).

Three-dimensional models for the trimeric structure of coronavirus spike protein were generated using AlphaFold2 (63) and the SwissModel server (64). Multiple sequence alignment in AlphaFold2 was performed using the mmseqs2_uniref_env and unpaired_paired parameters and the alphafold_multimer_v3 model. The remaining parameters were set to default.

Electrostatic potentials for the NTD regions of HCoV-HKU1 and other coronavirus models were calculated using the linearized Poisson-Boltzmann equation using the APBS software (65). PDB2PQR (66) was used to generate the PQR formatted files from the PDB coordinates by assigning partial atomic charges for the proteins using the PARSE forcefield at pH 7.0 and the solvent described as a dielectric constant of 78 and saline concentration of 0.15 M. The low dielectric cavity was set to 4.

## Results

Seventeen bat species were sampled at 12 sites (11 caves and one urban area), totalling 452 individuals and 712 biological samples processed (**Figure 1** and **Supplementary Table 1**).

We obtained two complete coronavirus genomes for samples from *Pteronotus gymnonotus* (Mormoopidae): Pgymn_N107_15 (30.414 bp) and Pgymn_N107_12 (30.423 bp) with average coverage depth of 621x and 314x (**Figure 2 and Supplementary Figure 1 and Table 2**). In addition, other seven partial genomes with a variable coverage breadth varying from 4 to 76% were obtained from the species *P. gymnonotus* and *Carollia perspicillata* (Phyllostomidae) (**Table 1, Supplementary Figure 2**). Full genomic annotation revealed a consistent gene order and gene length of the *Betacoronavirus* genus (**Figure 2A, Supplementary Figure 3**). The predicted spike protein length ranged from 1449 to 1453 amino acids and showed a more variable S1 and conserved S2 subunits (**Figure 2B**). Conserved domain analysis identified the N-terminal domain (NTD) within the S1 region including the trimer interface, but no receptor-binding domain (RBD) signature was detected (**Figure 2B**). Notably, despite the presence of the NTD, no clear amino acid alignment of the S1 region could be established when compared to homologs retrieved via BLASTp against the NCBI nr database, including well-characterized reference spike proteins from SARS-CoV, SARS-CoV-2 and MERS-CoV) (**Figure 2C** and **Supplementary File 1**). In contrast, conserved domains within the S2 region were identified, including the S1/S2 cleavage region, S2 cleavage site with the internal fusion peptide and the Heptad Repeat domains HR1 and HR2 (**Figure 2B and D**). Interestingly, the two complete coronavirus genomes were nearly identical except by the spike protein. Considering the 1449-1543 predicted amino acids, the pairwise protein similarity is ~79%, meaning that 1147 amino acid residues are identical between the two spike proteins. The majority of distinct spike amino acids are at the more variable S1 region that encompass the NTD and CTD regions (**Supplementary File 2**).

**Figure 2.**
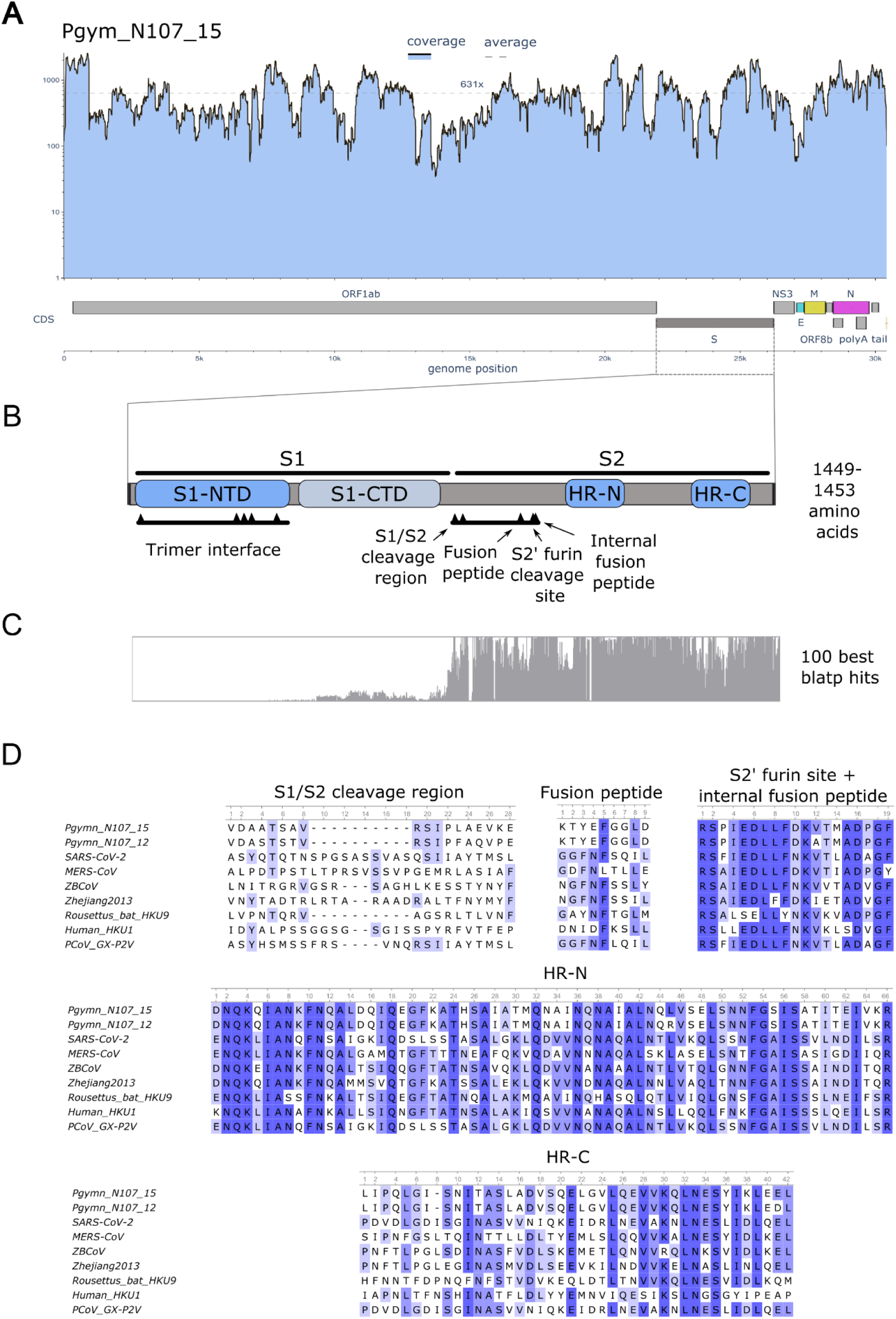
Genomic features and spike protein domain annotation of the *Ambecovirus* full genome recovered in this study. **A** - Coverage breadth, depth and complete genomic annotation of the new betacoronavirus genome; **B** - Schematic of the spike protein highlighting the S1 and S2 subunits, including conserved domains; **C** - Amino acid alignment of the *Ambecovirus* spike protein against the top 100 BLASTp hits, illustrating divergence in the S1 region; **D** - Amino acid alignment of the S2 domains found in the CDD search including the spike protein of the two complete genomes obtained in this study and reference coronavirus genomes.

**Table 1.**
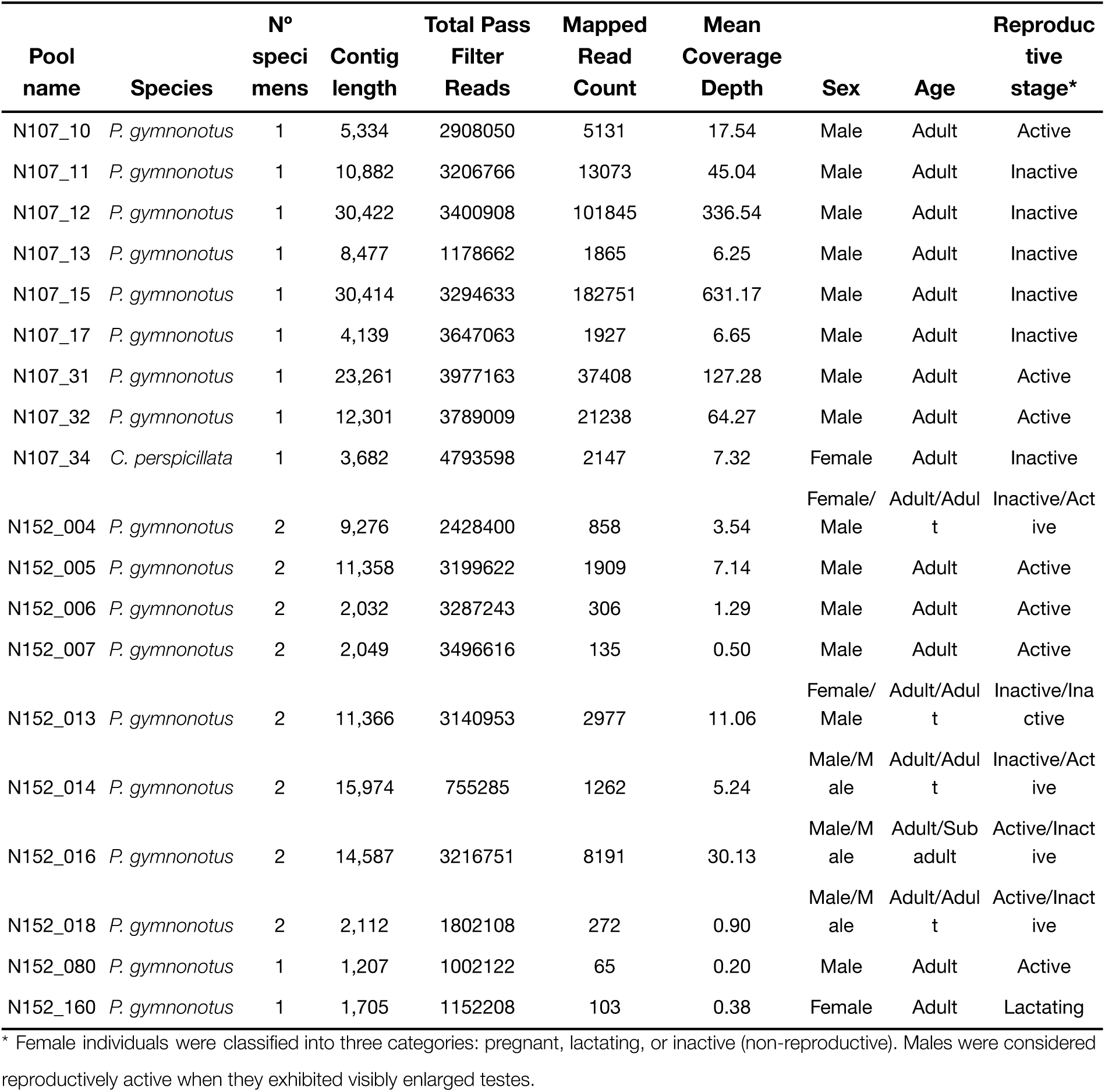
Descriptive statistics of complete and draft coronavirus genomes recovered in this study and metadata associated with the samples.

Structural modeling of the spike protein proved to be challenging, especially due to low sequence identity at the S1 region. None of the methods used reached acceptable quality metrics. AlphaFold3 yielded a pTM score of only 0.3, while SwissModel template-based strategy performed marginally better, achieving a GQME of 0.45 using the PDB ID 9BSW (67) of HCoV-HKU1 (**Figure 3A**). Nevertheless, a common feature among the models was the prediction of an NTD β-sheet motif. SwissModel template matching showed nearly 30% identity to lineage A betacoronavirus HCoV-HKU1 and HCov-OC43 for Pgymn_S15 and Pgymn_S12, respectively, two members of the *Embecovirus* subgenus within the *Betacoronavirus* genus. Both viruses use a β-sheet structured shallow groove in the NTD to bind to sialic acids containing-receptors. These residues are located within the first 210 residues of the S1 subunit for the HCoV-HKU1 and HCoV-OC43. A three-dimensional superposition of the predicted NTD structure of Pgymn_S15 by SwissModel and PDB ID 9BSW of HCoV-HKU1 is shown in **Figure 3B**. The β-sheet motif, which is also predicted for Pgymn_S12, is three-dimensionally well aligned suggesting ambecov S1-NTD would be contained within the same residue range. Prediction of the CTD was contrasting and poorly structured in all cases, as shown for Pgymn_S15 in **Figure 3C**. Although the CTD core secondary structure is predicted, there is a massive three-dimensional misalignment outside of the CTD core region. Based on this rather poor structural comparison to HCoV-HKU1 and HCov-OC43, ambecov’s CTD would likely lie around residues 310 to 600 (**Figure 3C**).

**Figure 3.**
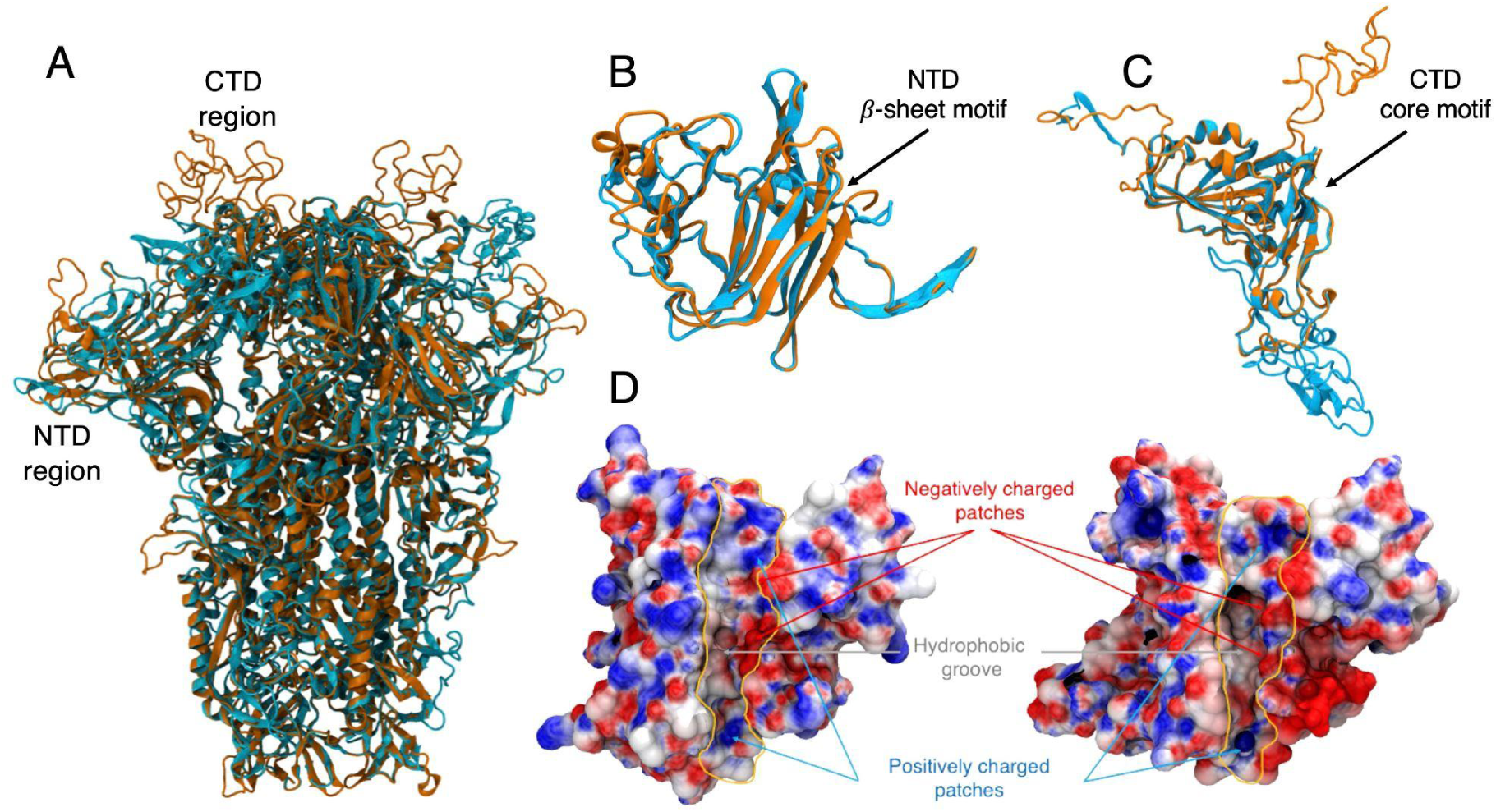
Superposition of a SwissModel predicted structure (orange) of the spike protein from ambecov based on the PDB ID 9BSW template from HCoV-HKU1 (cyan) depicted in cartoon model, and comparison of the electrostatic surface potential for the NTD region. **A** shows superposition for the spike protein in its trimeric form. **B** shows the superposition for the NTDs highlighting the predicted β-sheet motif for the model (orange). **C** shows the superposition for the predicted CTD region. The putative CTD region for the model (orange) is shown as highly unstructured, the result of a massive three-dimensional misalignment outside of the core region due to a low sequence identity between template and model. **D** shows similarity in the electrostatic surface potential for sialic acid binding β-sheet groove in the NTD from HCoV-HKU1 (left) and the corresponding region in the ambecov model (right) encompassed within thin orange lines. (Electrostatic potential ranges from −5kT/*e* (red) to +5 kT/*e* (blue) and it is plotted onto the van der Waals surface).

Another interesting fact about the spike protein from ambecov is that its total net charge at pH 7 is predicted to be equal to −30 *e*, the same charge adopted by HCoV-HKU1’s spike protein. It is worth noting that it dramatically contrasts to the charge of spike proteins from SARS-CoV-2, which tends to be around −6 *e*. Electrostatic properties of coronaviruses proteins have been well documented to play a major role in receptor, furin and RNA recognition, as well as immune evasion (68–71). Given the net charge similarity of ambecov and HCoV-HKU1 and its reasonably well-aligned NTD region, we have calculated the electrostatic surface potential for this region in both molecules. **Figure 3D** shows a similar pattern in the sialic acid β-sheet groove for the HCoV-HKU1 and the corresponding region in the model, supporting a similar functional role.

Phylogenetic reconstruction based on conserved *Nidovirales* proteins domains (3CLpro, NiRAN, RdRP, ZBD and HEL1 domains), including representatives of the four recognized genera (Beta, Alpha, Gamma and Deltacoronaviruses) of the Orthocoronaviridae subfamily placed the newly discovered genomes within the *Betacoronavirus* genus (**Figure 4A**). Moreover, these genomes clustered as a new, distinct clade, basal to the Sarbeco, Hibeco and Nobecovirus subgenera, having relatively high node support of 71.7 (aLRT), 83% (UFboot) and 0.99 (posterior probability) (**Figure 4A**). Together with a previously reported draft genome (BetaCoV_UNIFESP_unmBSS) the new genomes clustered with high node support (aLRT and UFboot = 100%) forming a novel subgenus we propose as *Ambecovirus* (American Betacoronavirus) (**Figure 4B**). Amino acid similarity thresholds within this subgenus at the 3CLpro, NiRAN, RdRP, ZBD and HEL1 domains ranged from 80 to 100%, while intergeneric similarity varied from 69% to 70% (**Supplementary Table 3**). Moreover, a more fine grained analysis of the nucleotide sequence revealed two main clades: Clade I (branch support = 99.3 aLRT and 100% UFboot) included all genomes generated in this study, a partial sequence obtained from *Pteronotus davyi* (Mexico, 2012), seven partial sequences from *Pteronotus parnellii* (Costa Rica, 2011-2016) and a basal sequence obtained from glossophagine bats (KX284064, Brazil, 2010) (**Figure 4B**); Clade II (branch support = 99.3 aLRT and 100% UFboot) comprised of partial sequences *Artibeus* spp. and *C*. *perspicillata* along with a draft genome (BetaCoV_UNIFESP_unmBSS) from *Artibeus lituratus* sampled in Ceará, Brazil, in 2023 (**Figure 4B**). This clade distinction was further supported by similarity analysis of the RdRp partial (346 bp). Clade I and II showed 69-100% and 96-100% intraclade nucleotide similarity respectively, while interclade similarity ranged between 69% and 75% (**Supplementary Table 4**).

**Figure 4.**
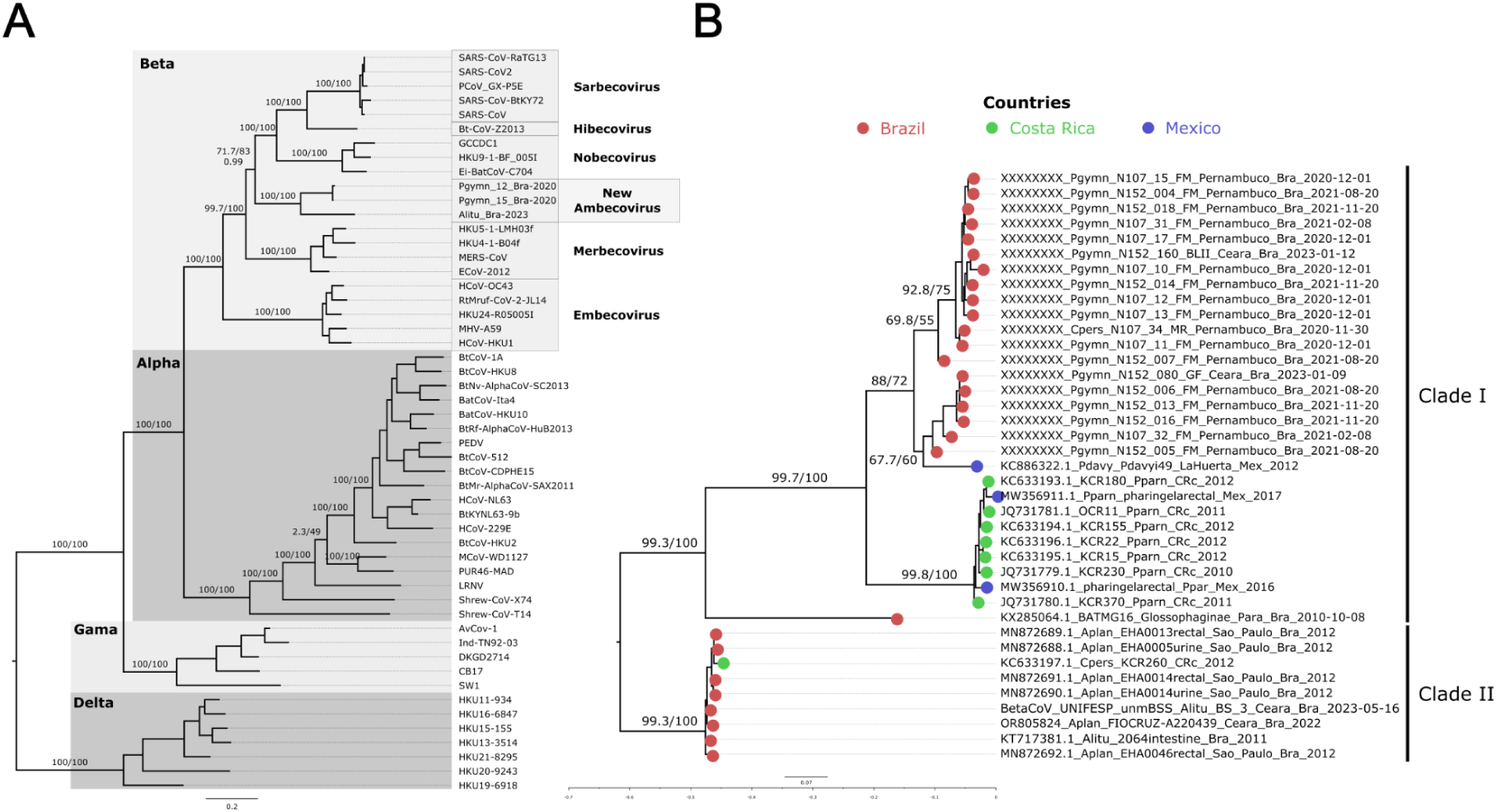
Phylogenetic reconstruction of the two complete coronavirus genomes characterized in this study and representative reference coronavirus genomes at the genus level. **A** - Maximum likelihood phylogenetic reconstruction based on amino acid sequences of *Nidovirales* conserved domains (3CLpro, NiRAN, RdRP, ZBD and HEL1 domains) - **Supplementary File 3**. **B** - Total evidence nucleotide tree based on the RdRp coding region including partial (439 bp) and full RdRp nucleotide coding region (~2790 bp) of the coronaviruses sequenced in this study - **Supplementary File 4**. Branch support values above nodes were calculated by aLRT/ultrafast bootstrap and posterior probability. Tip colors denote the country of sampling.

The sequences generated in this study originate from *P. gymnonotus* sampled multiple times in the same cave (Furna do Morcego, in Pernambuco state) from December 2020 to November 2021, but also from two other caves (Gruta do Farias, and Boqueirão das Lavras II) both located in Ceará state, where the same species was found infected. These two caves are located ~270 km distant from the caves in Pernambuco. Moreover, the *Ambecovirus* sequence was detected in November 2020, in *C. perspicillata* at the Meu Rei cave, also in Pernambuco, and less than 15 km from Furna do Morcego cave (**Figure 5**).

**Figure 5.**
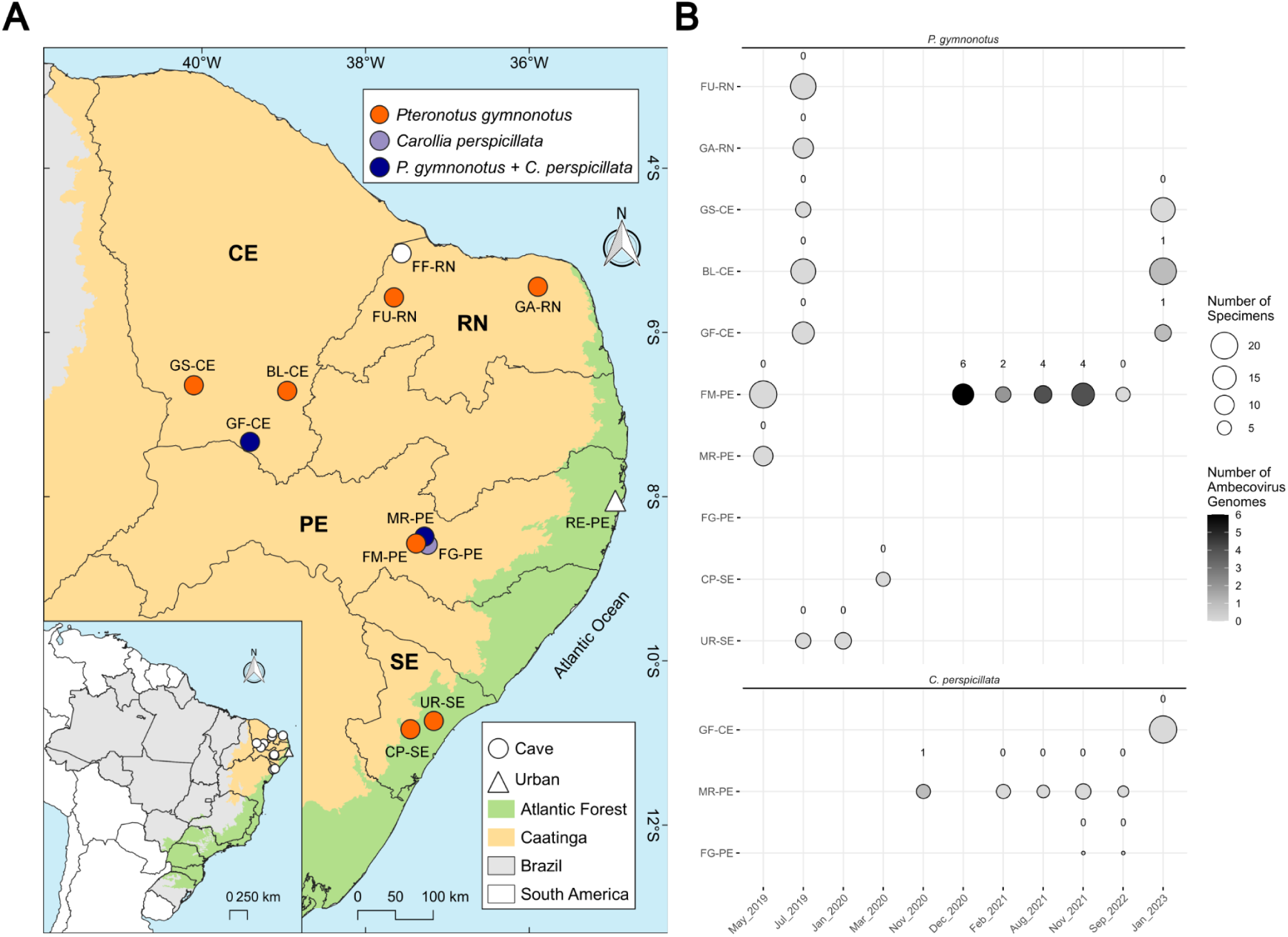
Spatio-temporal distribution of ambecovirus-positive samples obtained from *P. gymnonotus* and *C. perspicillata*. **A** - Map of the caves and urban sampling sites visited along the study showing colored caves where *P. gymnonotus, C. perspicillata* or both species co-occur. **B** - Number of individual samples per cave during the study period, including the number of partial or complete genomes recovered from metagenomic data.

## Discussion

Bats are known hosts of important zoonotic pathogens including coronaviruses, rhabdoviruses (rabies), paramyxoviruses (Nipah and Hendra) and filoviruses (Ebola and Marburg) (3). Despite that, efforts to characterize bat viromes are heterogeneous with highly biodiverse areas still understudied. Among these areas, the megadiverse biomes of the Neotropics, and particularly of South America, stands out: there, almost half of all known bat species were never screened for the presence of viruses. Moreover, the importance and impact of the rabies virus on the regional herds and on human health contributed to a clear bias in favour of its known host reservoirs (19). Additionally, the overwhelming majority of investigated bat species were screened using low throughput molecular techniques that rely on *a priori* knowledge and hence are able to detect a low viral diversity (3,19). To address these gaps, we conducted viral particle enrichment and meta-transcriptomic sequencing of cave and non-cave bats from sites in two Brazilian biomes (the Atlantic forest and the Caatinga drylands). This approach led us to the discovery of a novel *Betacoronavirus* subgenus, named *Ambecovirus*, circulating in multiple bat species across Brazil and the Americas.

The majority of zoonotic coronavirus lineages belong to *Beta-* and *Alphacoronavirus* (16). Betacoronaviruses are grouped into five known subgenera, from early diverging *Embecovirus* to more recent diverging ones namely, *Merbecovirus*, *Nobecovirus*, *Hibecovirus* and *Sabercovirus*. Embecoviruses have rodents as main reservoirs, while bats are the main reservoirs of the other four known subgenera (17,72). *Merbeco* (Vespertilionidae-specific) and *sarbecoviruses* (Rhinolophidae-specific) are known to spillover to other species, driving local outbreaks to large epidemics (72) suggesting viruses from these subgenuses are more prone to cross-species transmission, while *nobeco-* and hibecoviruses have been so far only detected in bats of the families Pteropodidae and Hipposideridae, respectively (73). Our phylogenetic reconstruction based on complete RdRp sequences revealed a monophyletic clade branching as a sister group of *Nobeco*-, *Hibeco*- and *Sarbecovirus* subgenera. Although partial Ambecovirus-like sequences have previously been reported in neotropical bats, their placement within the *Betacoronavirus* genus remained uncertain (18,33,40,41). Only one recent study (2025) recovered a draft ambecovirus-like genome so far (41). The position of ambecoviruses within the *Betacoronavirus* genus, and their detection being limited so far to neotropical bats (Mormoopidae and Phyllostomidae), implies long-term host-virus coevolution and regional isolation, consistent with broader studies that pointed out that Merbecoviruses and Alphacoronaviruses in the Americas are phylogenetically distinct and less prone to cross-species transmission than from their Old World counterparts (18). More broadly, the current data suggests that an ancient split occurred in the *Betacoronavirus* genus separating ambeco-from nobeco-, hibeco- and sarbecoviruses in the Americas while these last three shared an Old World common bat ancestor. Interestingly, this finding further corroborates other broad scale studies on coronavirus evolutionary history which showed that merbecoviruses and alphacoronaviruses infecting bats in the Americas are isolated from the African and Asian coronaviruses (18). Our data supports regional variation in the long-term evolution, but at a more recent time we detected one potential host-switching event between distantly related species (from two different neotropical bat families - Mormoopidae and Phyllostomidae) or this detection in multiple bats species suggests that ambecoviruses have broad host range infecting neotropical bats. Further studies are warranted to evaluate these hypotheses.

### Evolution and zoonotic potential

The detection of a new *Betacoronavirus* subgenus raises a number of questions about its evolution and zoonotic potential. The full genomic sequence recovered in this study allows us to perform an in depth genomic annotation and characterization of the predicted proteins to test some hypotheses. Betacoronaviruses differ in the composition and order of genes at the 3’ terminal region, after the spike protein. The envelope, matrix and nucleoprotein are located downstream to each other followed by hypothetical proteins in *Nobecovirus* and *Merbecovirus,* while envelope and matrix are in tandem and separated from the nucleoprotein by two to five hypothetical proteins in *Sarbecovirus* and *Hibecovirus* (https://ictv.global/report/chapter/coronaviridae/coronaviridae/betacoronavirus). The *Ambecovirus* subgenus revealed a synteny similar to *Nobecovirus* suggesting conservation of the gene order of the ancestral lineage (**Supplementary Figure 3**). Using available protein conserved domains and structural information for a number of medically important Betacoronaviruses, we characterized in more detail the spike protein responsible for receptor binding and a key molecule responsible for the infection capacity of coronaviruses. The spike protein is divided into two subunits, the more variable S1 region and the more conserved S2 one. S1 is further divided into S1-NTD and S1-CTD, while the second is mainly responsible for the fusion of the viral particle with the host cell membrane in merbeco and sarbecoviruses, the first can also bear the receptor binding bind domain to sialic acid receptors in embecovirus (74). The S1-CTD contains the receptor-binding domain and mediates receptor interaction in Merbecovirus and Sarbecovirus, but it also plays an ancillary role in Embecovirus by stabilizing the spike trimer and modulating the conformational transition between pre-fusion and post-fusion states (74). These two regions are also the main targets of neutralizing antibodies and because of the strong diversifying selection pressure, they normally evolve faster than the S2 region (75). Interestingly, *Ambecovirus* spike proteins characterized in this study showed a NTD identifiable conserved domain but lacked an identifiable RBD domain and RBM motif. Moreover, there is a complete lack of alignment of the spike S1 region with any other spike proteins available in the databases, including those of the well characterized human pathogens (MERS-CoV and SARS-CoV-2). Such pattern was already observed between the S1 subunit of different Orthocoronaviridae genera that share little sequence similarity, yet it departs from the generally accepted view that congeners share higher sequence similarity as reported by Li (75). Therefore, the NTD and RDB region of the *Ambecovirus* are highly divergent at the amino acid level.

In order to further investigate if structural homology still exists between the S1 NTD and RBD region among Ambecov and the other Betacoronavirus subgenuses we sought to perform 3D modelling of the entire spike protein recovered. While the S1-NTD 3D modeling is compatible with a β-sheet shallow groove also found in other two Embecovirus spike 3D structures, the S1-CTD showed distinct conformations at every modelling attempt. In addition, ambecov’s CTD does not contain sequence equivalence to the conserved residues found in ACE2 binding sarbecoviruses, such as Y449, G496, N/Y501 and Y505. Unlike sarbecoviruses, the CTD in HCoV-HKU1 and HCoV-OC43 plays ancillary roles, such as structural stabilization of the trimeric structure, modulation of pre-fusion to post-fusion conformational transitions, and immune evasion through glycan shielding (16,74). Hence, it is possible that the spike protein interaction with the receptor in these bats either occurs through different contact regions and/or uses sialic acid cell receptors. Another important implication is that, due to high sequence divergence, the ambecoviruses may not be able to infect human cells. However, both hypotheses must be rigorously tested using *in vitro* and *in vivo* assays to precisely evaluate its zoonotic potential. Once ambecoviruses infects both sylvatic and synanthropic bats and cohabitation with other hosts including humans is a premise for virus cross-species transmission, a more in depth investigation is needed to understand the infection capacity of these viruses in other animal cells, as well as to track and reveal its ecological dynamics in the wild, including biotic and abiotic factors that drive viral shedding and potential spillover events.

In addition to the deep ancestral inferences, high resolution phylogenetic reconstructions also allowed us to make inferences about the more recent *Ambecovirus* evolution. There are at least two clades circulating in the Americas: clade I, in sylvatic and sinantropic bats (*Pteronotus* spp. and *C*. *perspicillata*); and clade II, which circulates mainly in sinantropic bats (*Artibeus* spp. and *C*. *perspicillata*). Our findings also suggests that clade I strains infects mainly *Pteronotus* spp., but may be able to naturally infect multiple sympatric host species, i.e. species roosting in the same cave (*P. gymnonotus and C. perspicillata*, for example), or able to cross species barriers. Habitat overlap, and particularly host density at roostings with multiple bat species, have been shown to facilitate multispecies infection by coronaviruses due to close physical contact (76). However, more longitudinal data (e.g. shedding) from *P. gymnonotus* and *C. perspicillata* is needed in order to assess if our results represent an isolated event leading to virus extinction or if sustained transmission among *C. perspicillata* populations has been established. We were able to sample the same bat metapopulations in several time periods, but most of our sampling was restricted to the dry seasons, hence the temporal virus dynamics captured is limited for broad inferences about infection prevalence and shedding across time. In case sustained transmission is confirmed, it shall have two major implications: a) it will indicate cross-species transmission of *Betacoronavirus* in neotropical bats, contrary to the reports of neotropical bats-*Betacoronavirus* cospeciation and the implied resistance to host switching (40); and b) the detection of the new ambecovirus in *C. perspicillata,* which is frequently found roosting in caves but is also a common synanthropic species, raises risk concerns, as this species is frequently found in large urban centers (77,78).

Interestingly, Anthony *et al*. (18) have pointed out that the absence of coronaviruses in some bats species was likely due to insufficient sampling. This is confirmed by several studies showing that only a third of the bat species were investigated for virus infections so far (73). In our study only *P. gymnonotus* had more than 200 specimens sampled, while an ideal sample size of more than 150-400 individuals is recommended for increasing the chances of detecting Coronaviruses in bats (18). However, population sizes for many of the sampled bat species is reduced with sometimes less than 25 individuals in a colony (i.e *Desmodus rotundus* at the Furna do Gato cave). Therefore, our sampling reflects relatively well the natural population sizes of these species. Still, the low sample size may have reduced the chance of detecting ambecoviruses in some of the studied bats. In addition, not only limited sampling contributes to such underestimation, but also the use of specific assays relying on previous information, particularly for highly divergent viruses such as ambecoviruses. Therefore, more frequent bat sampling and viral screening with unbiased methods are necessary to further characterize new coronaviruses and other zoonotic pathogens.

### Perspectives

The discovery of the *Betacoronavirus* subgenus-*Ambecovirus* in neotropical bats, besides bringing new insights to viral evolution and ecology of these bats and viruses, raises some important questions: Are bats the main reservoir of this new subgenus? Does *Ambecovirus* infect only bats or also has the potential to infect other mammals, including humans? How frequently do these viruses cross the species barrier? While current evidence suggests host restriction to bat species (18,28,40) for Alphacoronaviruses and Betacoronaviruses of the *Merbecovirus* subgenus in the Americas, the detection of *Ambecovirus* in both sylvatic and synanthropic bat species raises concerns about potential cross-species transmission. Further understanding of the host range and specificity at molecular level also will strengthen our understanding of spike protein evolution, particularly of the S1-CDT region that might contain the RBD and RBM motifs. For this new CoV subgenus open questions also remain regarding the receptor used and the domain and residues that are more important for binding to the cell receptor ultimately defining tissue trophism, host range and potential epidemic potential. The recombination potential of Ambecoviruses also needs to be addressed in the future. Recombination is a well-documented driver of coronavirus evolution and emergence, as seen in SARS-CoV and MERS-CoV. However, due to the lack of closely related parental lineages and limited genomic data, recombination analysis for Ambecoviruses remains inconclusive. As more complete genomes are sequenced, it will be possible to assess recombination frequency and identify potential hotspots. Lastly, at the population level, there is a need to further assess viral shedding and map the likelihood of host overlap with other wild and domestic species or humans to more accurately estimate the spillover and emergence risks. In summary, *Ambecovirus* represents a deeply divergent lineage within *Betacoronavirus* genus, and its discovery underscores the importance of continued virome exploration in under-sampled regions. Addressing these open questions will not only clarify the evolutionary trajectory of this subgenus but also inform risk assessments for future zoonotic events.

## Data Availability

Supplementary materials are available at - Wallau, Gabriel (2025). Ambecovirus, a novel Betacoronavirus subgenus circulating in neotropical bats sheds new light on bat-borne coronaviruses evolution. figshare. Dataset. https://doi.org/10.6084/m9.figshare.29602244.v1.

## Acknowledgements

We are very grateful to all the people who helped us during the fieldwork, especially Bárbara Coelho, Deibson Belo, Eduardo Pires, Jennifer Barros, Juliana Bezerra, Maria Júlia de França, and Narjara Pimentel. We also thank Heike Baum, Alexandra Bialonski, and Marike Petersen for their excellent technical assistance in metagenomic sequencing (mNGS).

