## Supplementary Figure 1 for "*Ambecovirus*, a novel *Betacoronavirus* subgenus circulating in neotropical bats sheds new light on bat-borne coronaviruses evolution"

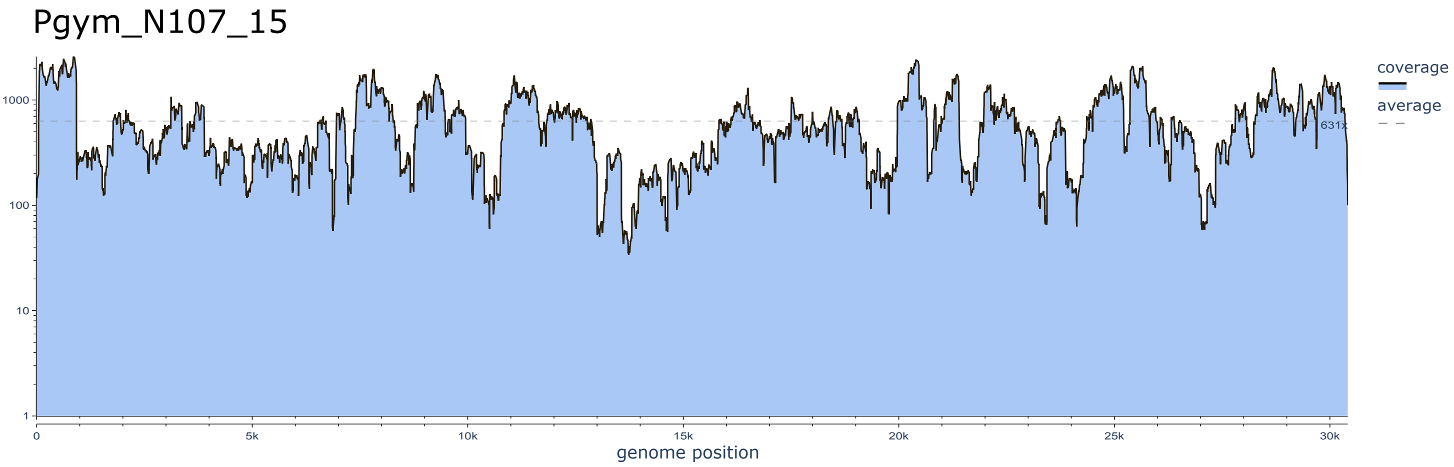
Pgym_N107_15
coverage
average
631x
1000
100
10
1
0
5k
10k
15k
20k
25k
30k
genome position

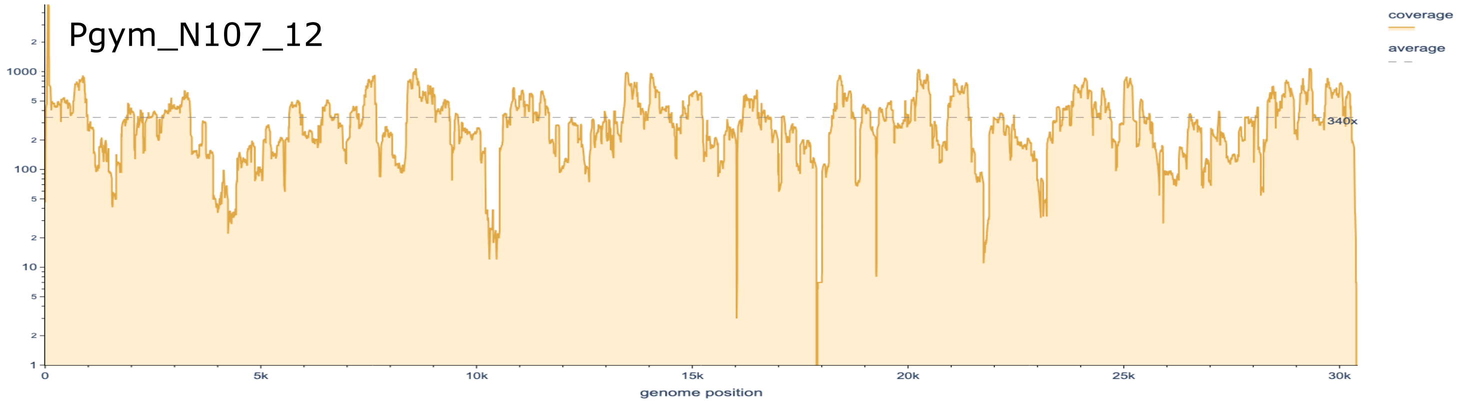
Pgym_N107_12
coverage
average
340x
1000
100
10
1
0
5k
10k
15k
20k
25k
30k
genome position

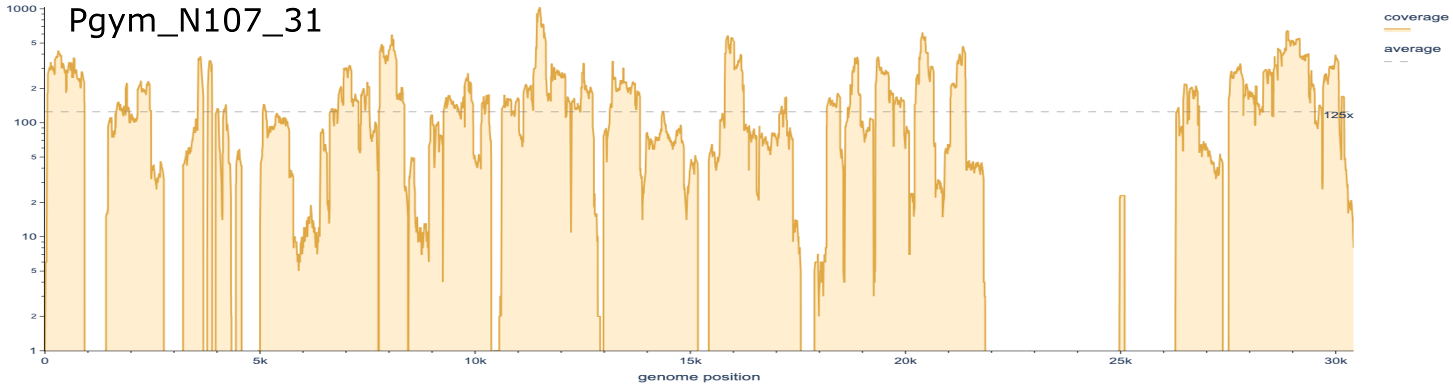
Pgym_N107_31
coverage
average
125x
1000
100
10
1
0
5k
10k
15k
20k
25k
30k
genome position

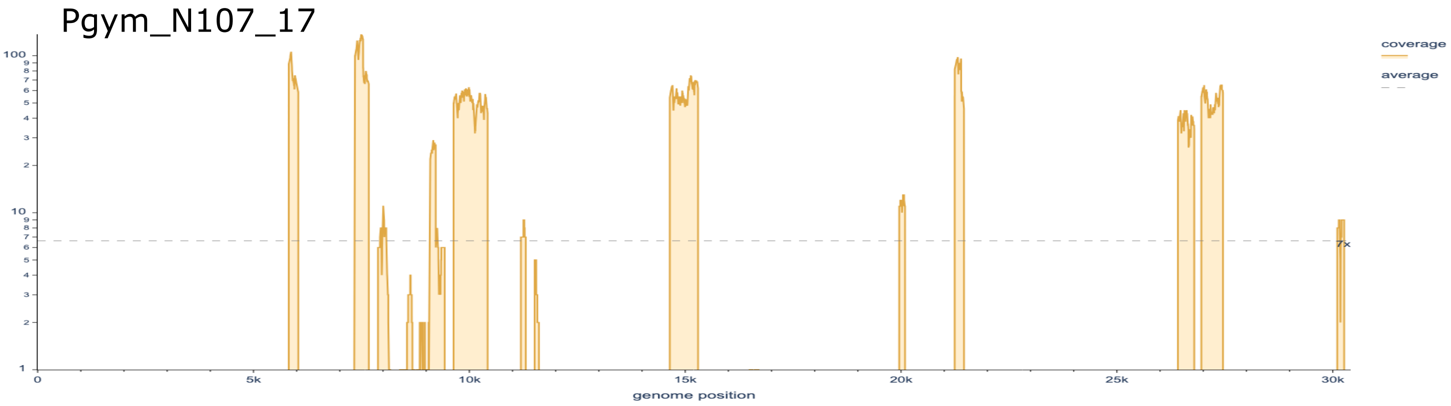
Pgym_N107_17
coverage
average
7x
100
10
1
0
5k
10k
15k
20k
25k
30k
genome position

Pgym_N107_10

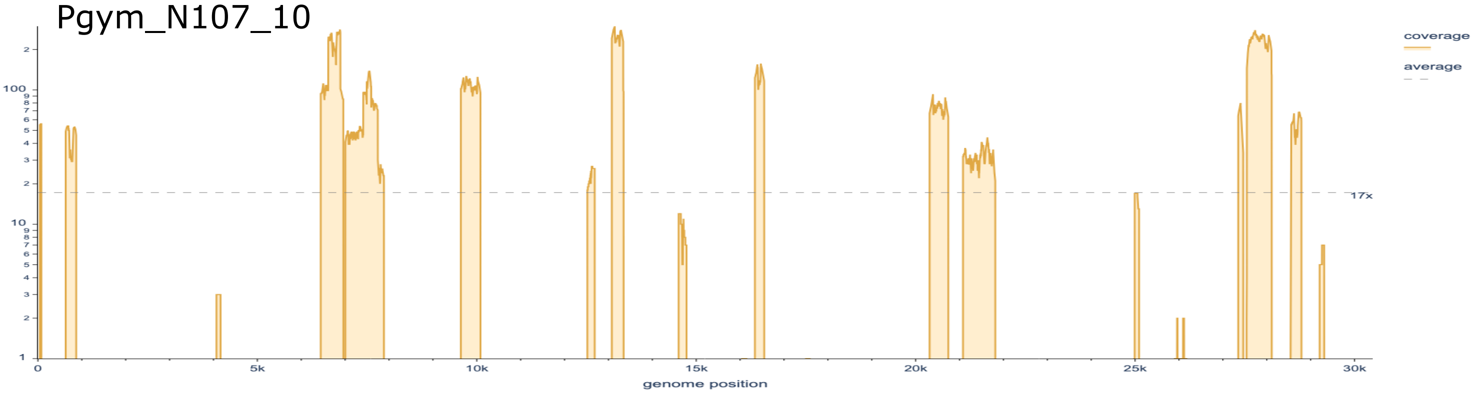


Pgym_N107_13

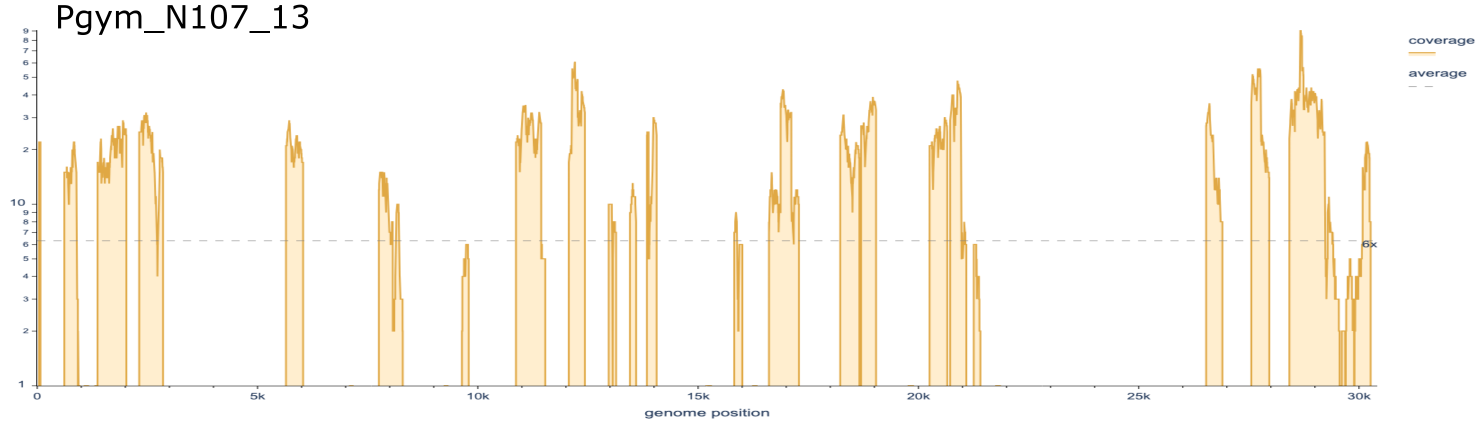


Pgym_N107_11

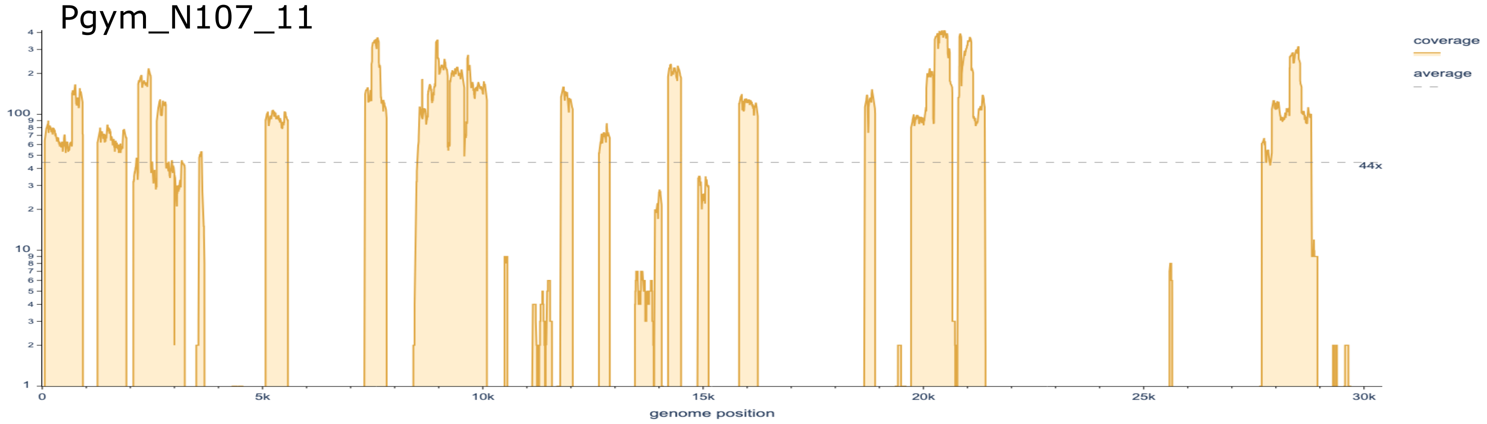


Cpers_N107_34

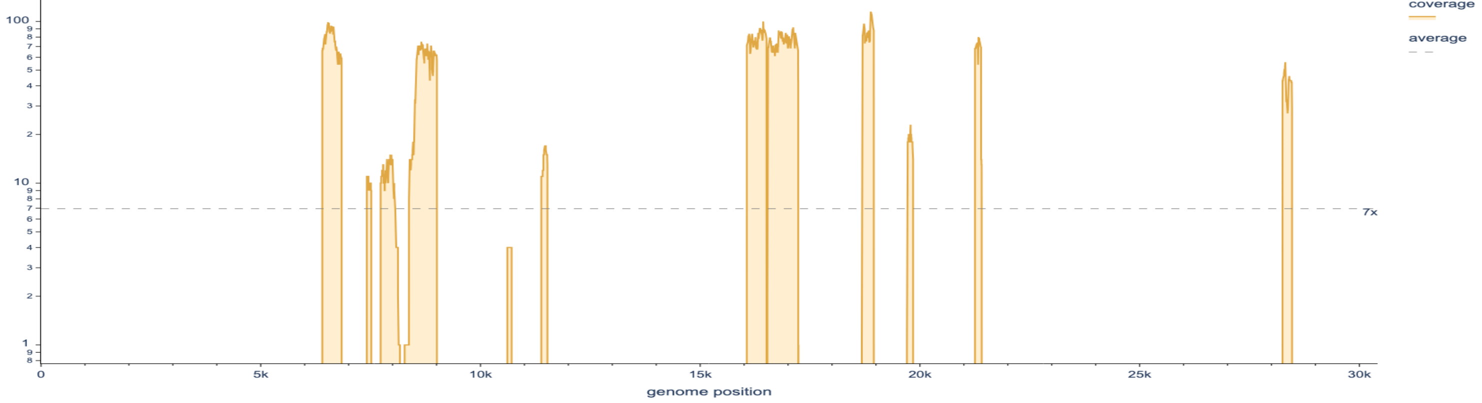

Pgym_N107_32

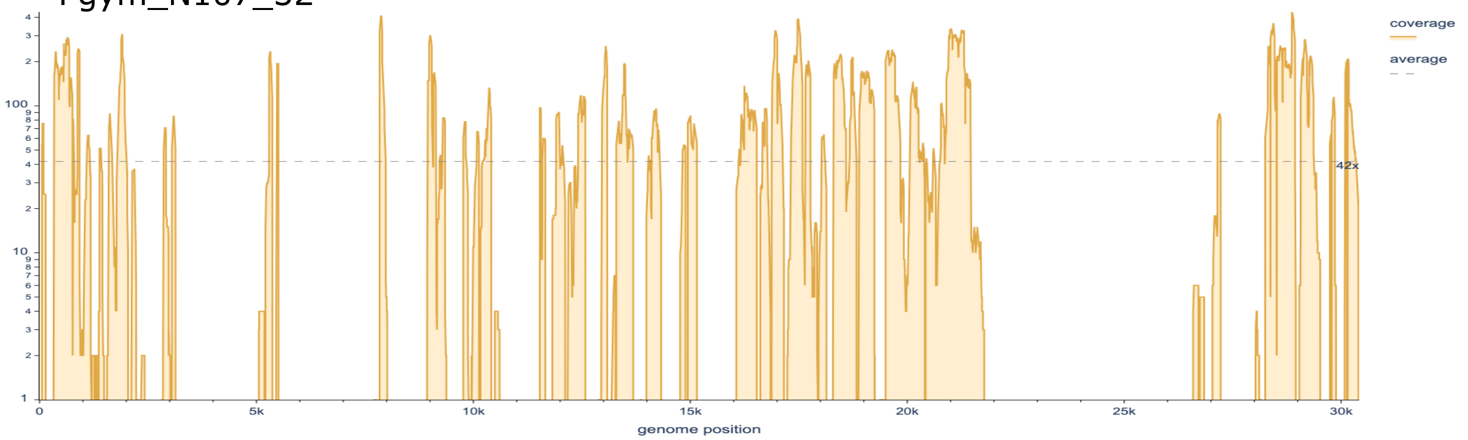


Pgym_N152_004

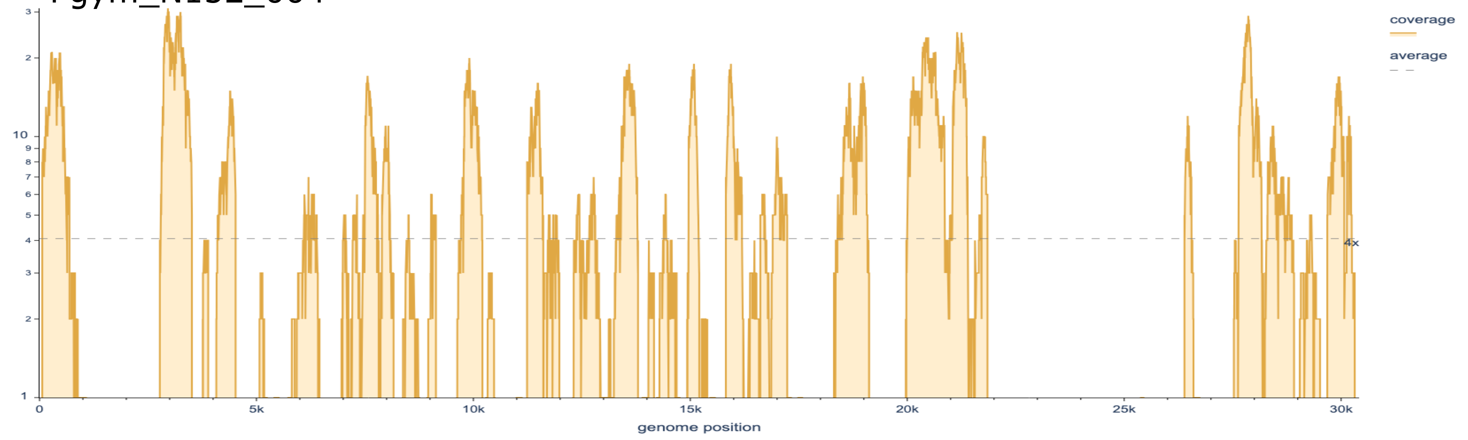


Pgym_N152_018

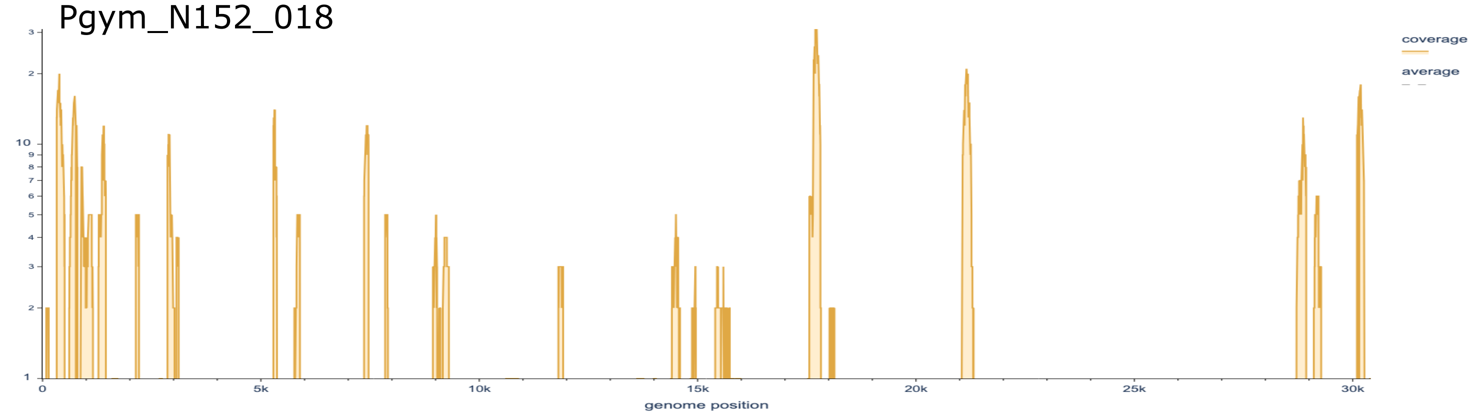


Pgym_N152_160

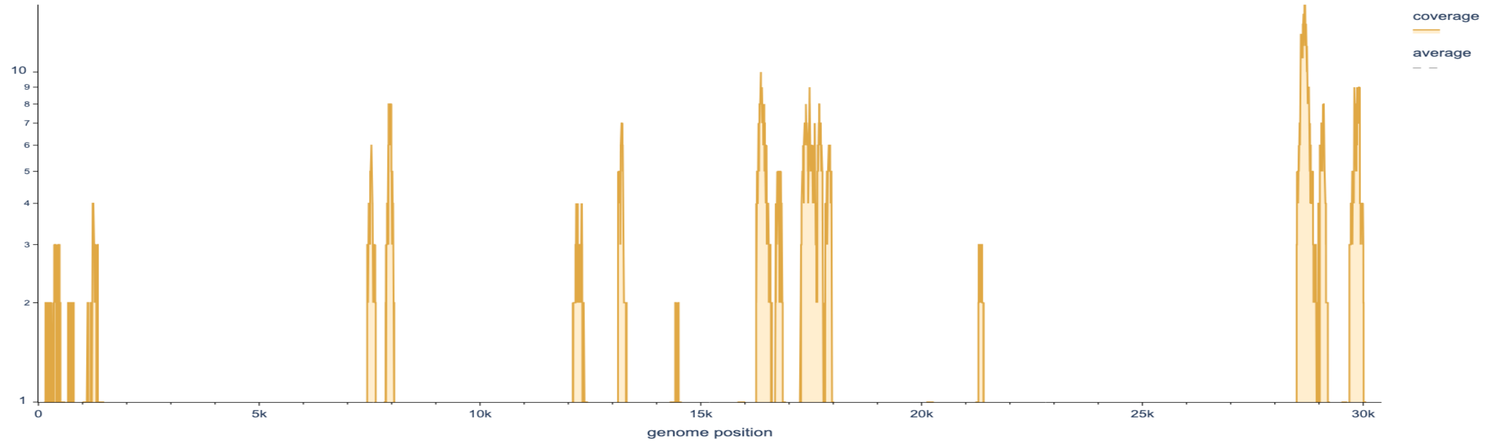

Pgym_N152_014

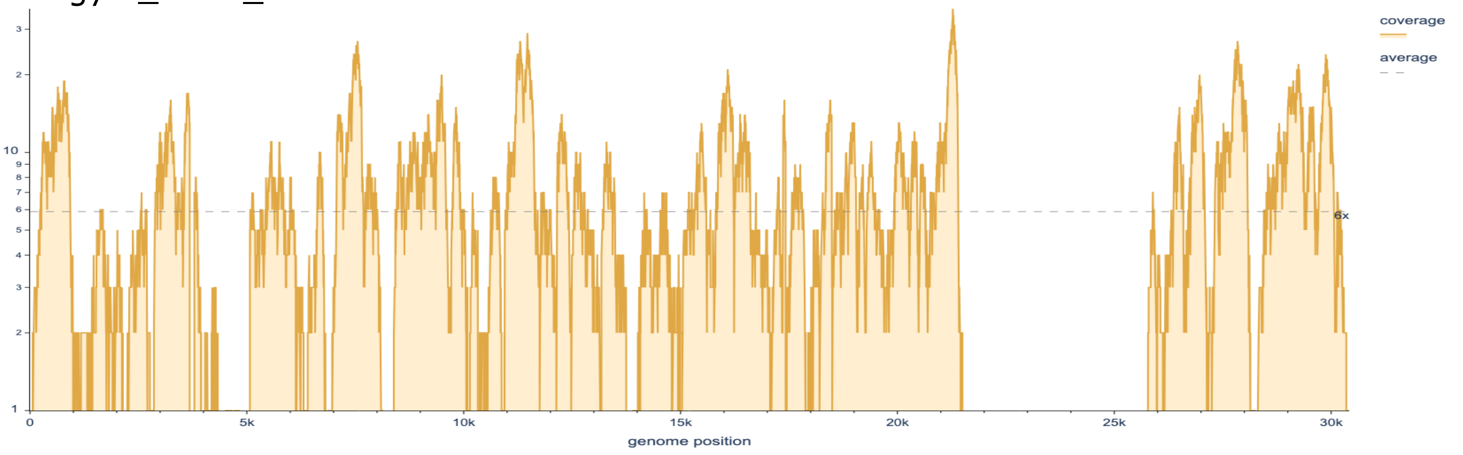


Pgym_N152_007

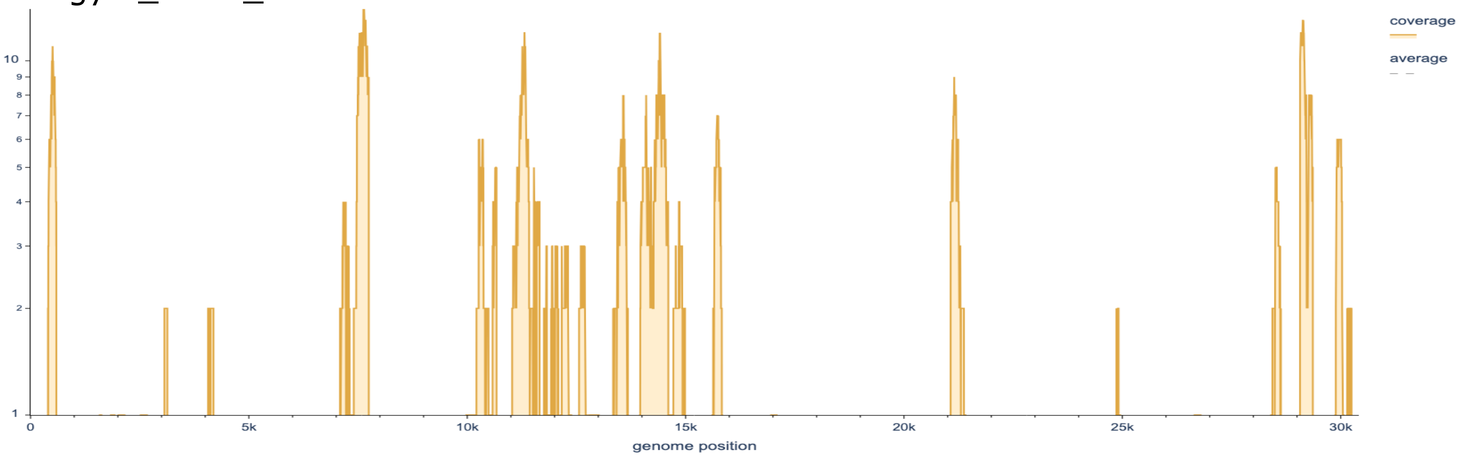


Pgym_N152_080

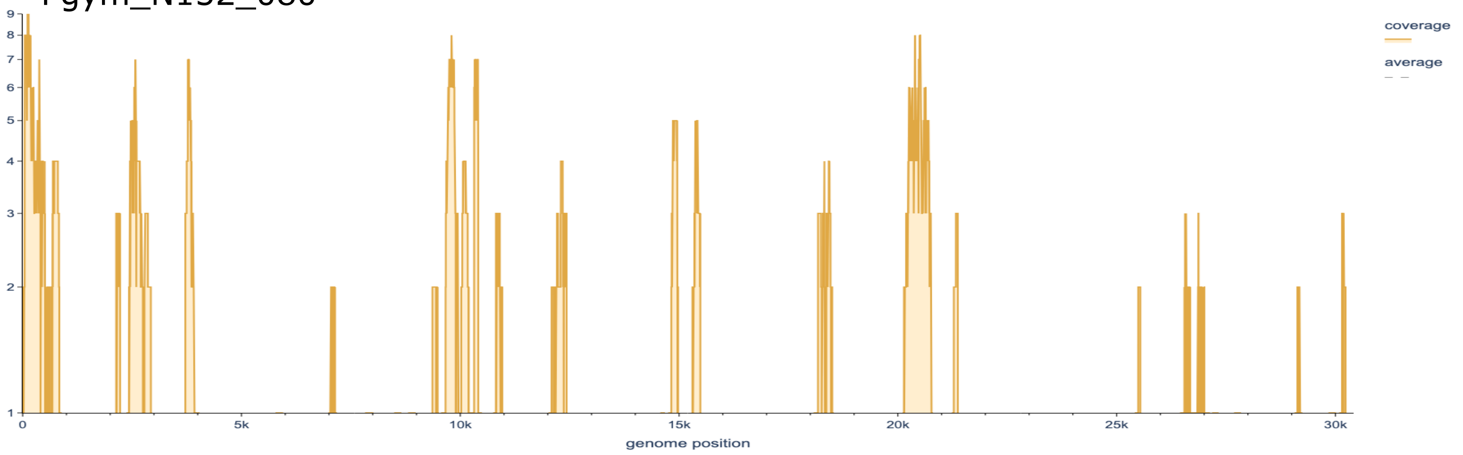


Pgym_N152_006

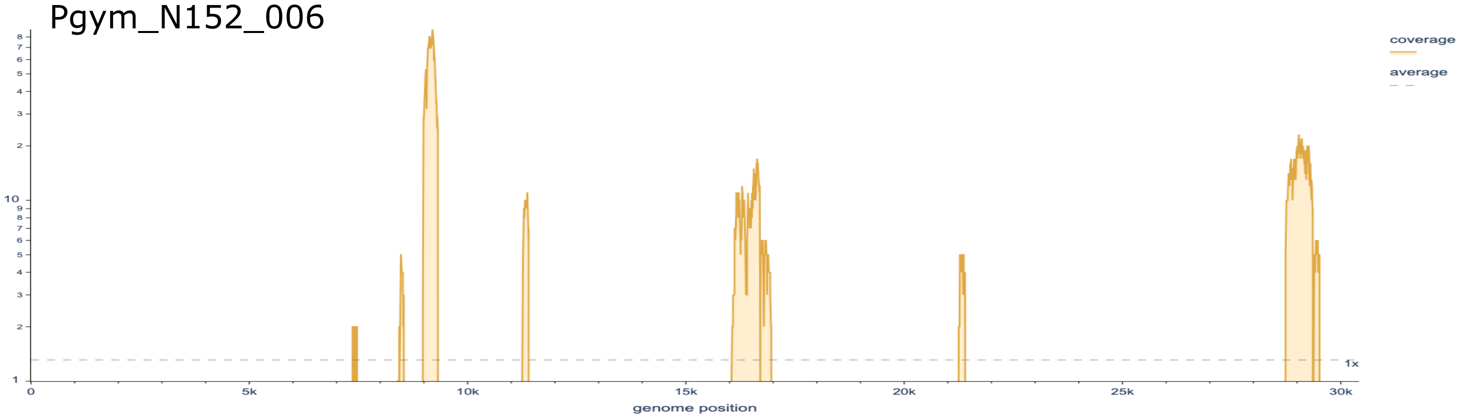

Pgym_N152_014

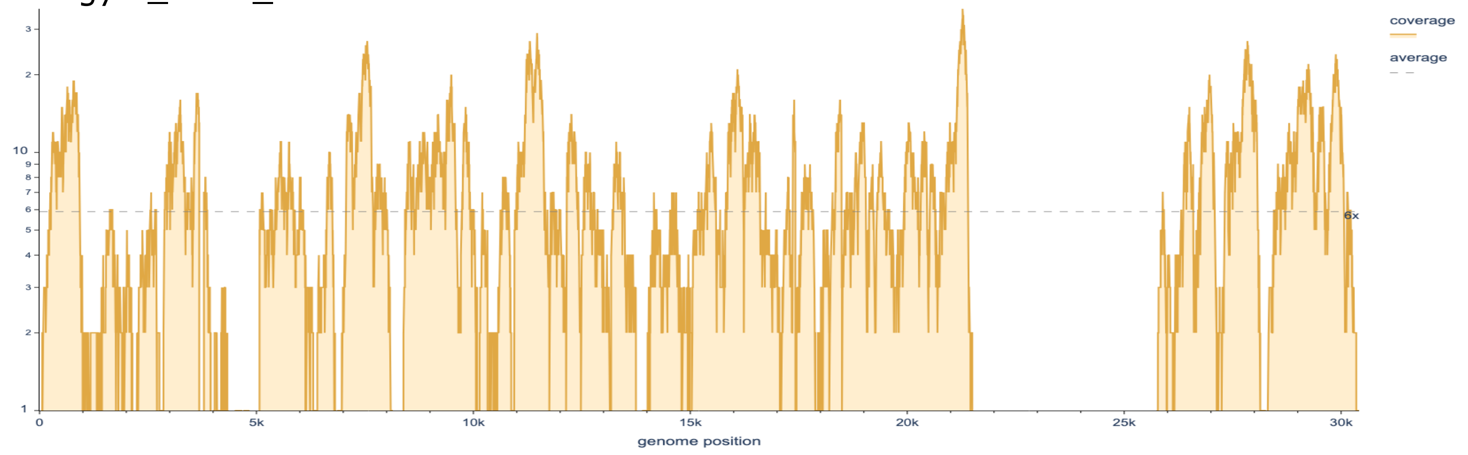


Pgym_N152_013

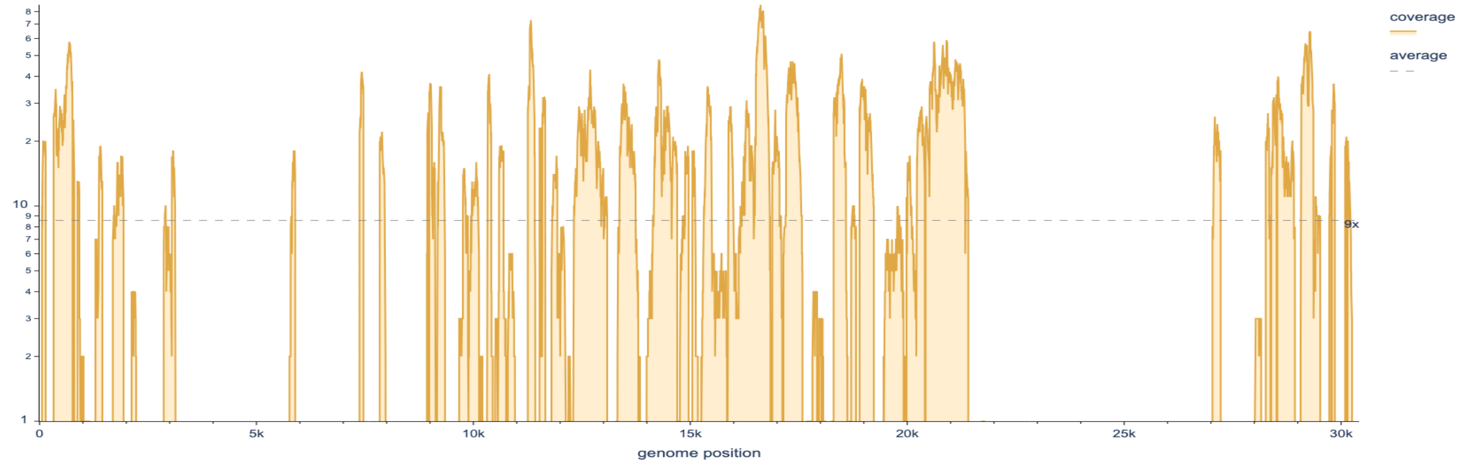


Pgym_N152_016

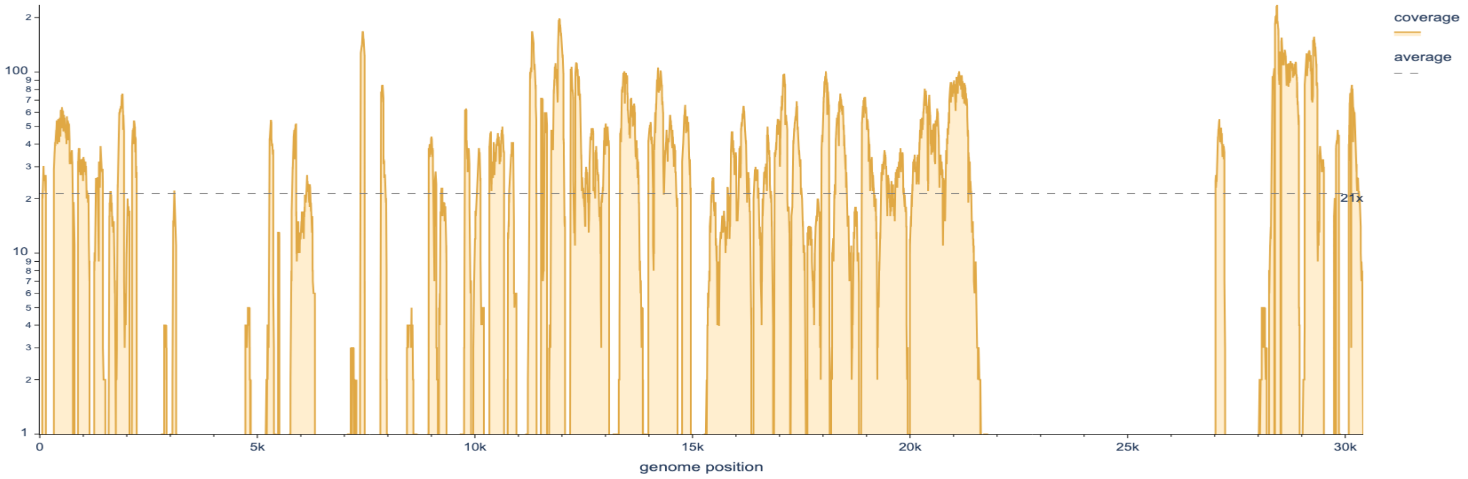


Pgym_N152_005

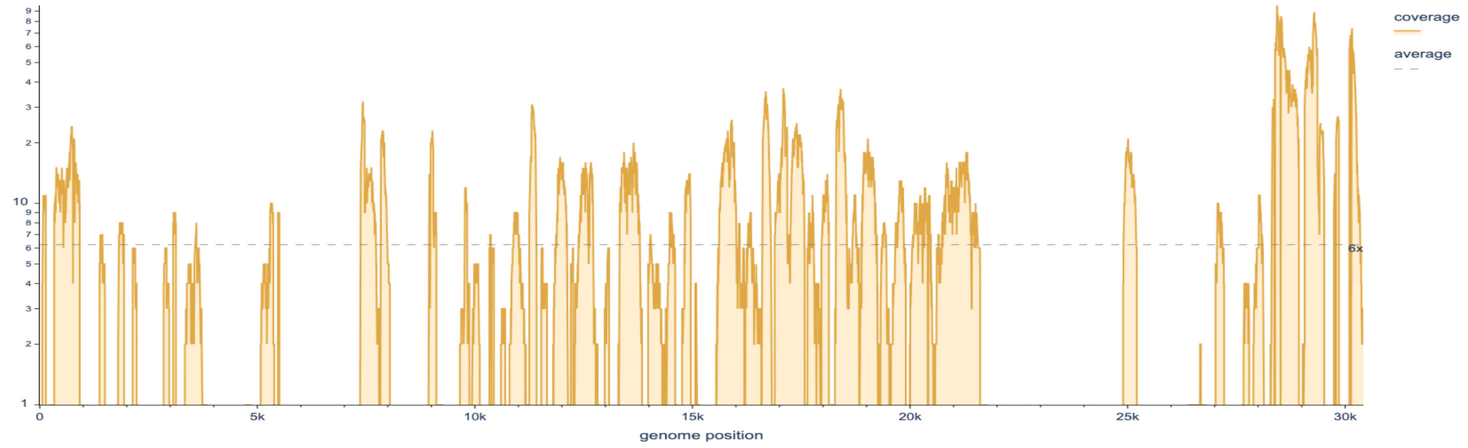
