## Supplementary figures and images for "*Ambecovirus*, a novel *Betacoronavirus* subgenus circulating in neotropical bats sheds new light on bat-borne coronaviruses evolution"

### Supplementary Figure 2

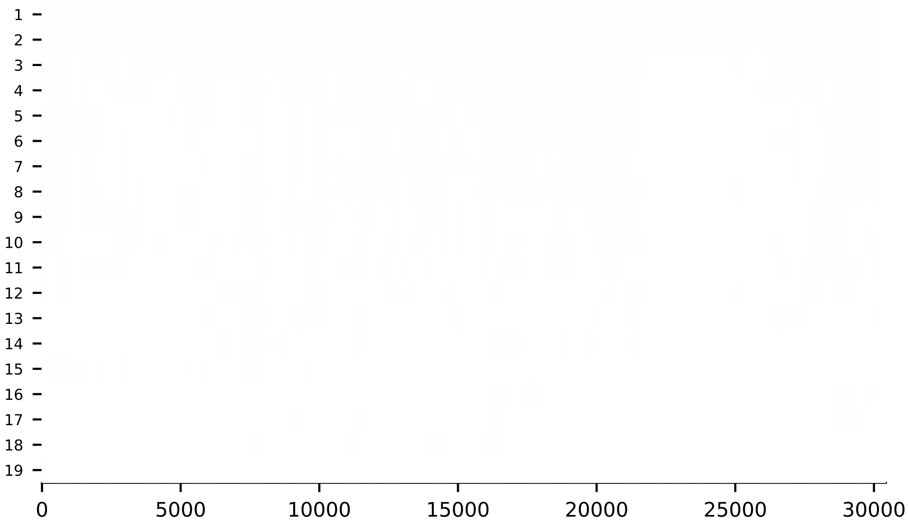

### Supplementary Figure 3

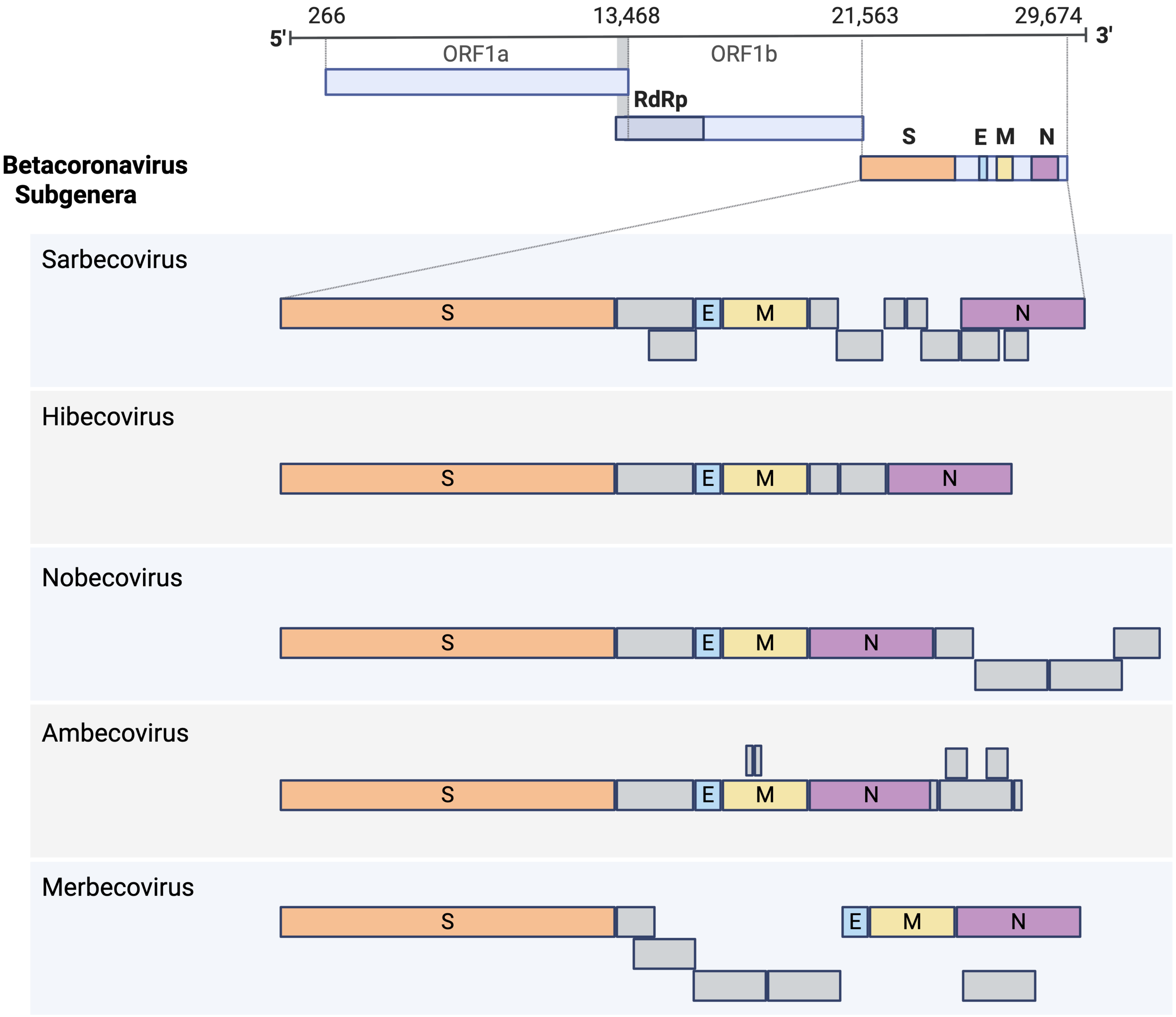
