## Supplementary File 2 for "*Ambecovirus*, a novel *Betacoronavirus* subgenus circulating in neotropical bats sheds new light on bat-borne coronaviruses evolution"

```
1 - 150
N107_S12  HRQFSGPMASLRTIVLLLCVHRAFSACSLKSDSDTACPNDAKLQEAAPLVANPFIDATKVSIAMKPLIYGTACYRRVTRTGRPNNIGSSPMTQLGVVYNVVDRMYTNTTTLVTQALPSAADGWKTITVAHRYSVCSSRTTPIRVMCVPDS
N107_S15  ------PSSSMAALILLLVVG-AYASCPLTSDNDASCPDNAVLQRSDPLVANPFIQQVSVSTAMVPLIYGTSCYRSIARSTRANPTSNQPIVEIGVVYNIVDKLLSNTTTTVTQPLPTLTAGWTVVPTAHKYSVCSSRTKGPRAMCVPTS

151 - 300
N107_S12  YSLPDNIDKWVPVSEGVTFSYAPTTNTHMIAPAVLSVPLGDSITGYSLYVGPTRWWPERGNDTIVILYCNFTMCPAAYSFANLGYWHHGENSYDRPLLSEQLFYGQSVHCLVNMTLIVPFGQDLVLSFIQDDAFVYGYAGSSKLSASILF
N107_S15  YTLPSSVDTFIPLTDGVVFKYAPQSNVHMFAPVILTVPMGDVVTGYSLYVGPTRWWPERGNDTITVMYCNFSMCPSAYVFANLGYWHHGEGSYDRYLLSEQLFHGSSVMCIVNTTITIPFGTDVVLSFVQDDAFLYGYVGTSSRSATVLF

301 - 450
N107_S12  RQPLQNKFSHFKFIPITYMSDSVQITAVGTYALLSKRTYLVRYDLQGMAEEYADCDGSELHQMLCNAGTFKPRDGVYKLRPVTDTPQLTVGPRVVRKETCDLNRLLSKDGAGKFKLNHPGFHVSKFYNCGKRSCGKPANNDVTNVVHDRA
N107_S15  RTPLQNKLSHFKFVPITYTSDNVQLISSGTYALLTRRNYLVQFDLQGFAVRFADCDGETLHQLFCTAGTFTPVDGVFRLKPLVDTPQLTVGPRVVSKAKCDLSRLISPDGPGKFKLNYPDFHVSKFFRCGKLACAKRTSTGVTDIQINLD

451 - 600
N107_S12  ACPSGSQPLMVQKV-CQNGTYADWLTPAIKSQTISGCEYDSAAVREAFKRMSVKCFGISSDMLFQSCFGSITYYHLPIHTGEKGHRMFDPTESNAFRKYGLGVDTGKPYCVAFGSMTVSSHSTFSYRDGPPNKLANCYLDLKPDCTLKS-
N107_S15  SCPPGYEELMVQGTGCKTDTYADWLTPAIRTQTISDCEYDSQAVREAFKRLSVKCHGISADSLFQSCFGSITYYHLSIHAGEKGHRMFEPTDSNAFRKYGLGVDTGRPYCVAFGAMNVRSHDLFRHSEGPSNKLANCYLDLKPDCTLKAG

601 - 750
N107_S12  GSGTYFECAGYKPGSTAIGNFDRNKLIAQHLPGRACGHVFGTHSGFSSGYAAYPTLPGQIPPYTAQPCGP-HAVGD-QPALGQIGFKPMQCDVAGGDGEWKGDFAIIMSLTLNSDSMVCPTKNALPSVFDHTSKCVTYDFGDISGQGVFL
N107_S15  GSQTYYECAGIWPGGTGVPLDKKKETLARKDSGHACGYKFGTHSGFKPDYKAYPAVPGQVPPYTSQPCGPLHADGTAAPTYGQAGYKPMLCDMA-QDDQWSGDFAIVMSLTLNSDSMVCPTKNVLPSVIEHTLKCVNYDFGDVSGQGVFL

751 - 900
N107_S12  KPGASYMGFVCKAPAFMGRYYVYDQLNHFVGYWYDNSTYLCVTPCKIVQREAIYQSKPHWAAGQLPILYRSAKCADVRASGIDTEVGCVTVNTNATAQFCSDMSTAIGASLCVDASTSTVRSIPFAQVPEADPIFKDSIFIATNYSFVSN
N107_S15  RTGQRYMGFTCRAPAFTGRYYVYDQLDHFVGYWYDNVTYLCVTPCKIVHRDAVYQRAPHRATNALPILFRSAKCADVSQNGVDTSVGCVTVNTNDTDKFCADMNTAIGASLCVDAATSAVRSIPLAEVKEADPIFKDSIFIATNYSFVSN

901 - 1 050
N107_S12  AQYPQTHAPKISIDCESYICSGHFRCLELLRQYGSFCVEINKALSGVALRQDRSAVQVLNAMRLSSQAGKTYEFGGLDIGNKFMIVAQGRSPIEDLLFDKATMADPGFMTDYQKCIEAKGAGLAAYEDADIQCVQKYNGVAVLPPIMSDG
N107_S15  AQYLQTHAPKISIDCESYICSGHFRCLELLKQYGSFCVEINKALSGVALRQDRSAVQVLNAMRLSSQAGKTYEFGGLDIGNKFMGVAQGRSPIEDLLFDKVTMADPGFMTDYQKCIEAKGAGLAAYEDADIQCVQKYNGVAVLPPIMSDG

1 051 - 1 200
N107_S12  LIGKYSAALGAGMQFNALTGGLLGYGMSATYTMAIAARINALGVTQDVLMDNQKQIANKFNQALDQIQEGFKATHSAIATMQNAINQNAIALNQRVSELSNNFGSISATITEIVKRLDQLESDVQIDRLINGRMTALNTYVTQTLLDIKR
N107_S15  LIGKYSAALGAGMQFNALTGGLLGYGMSATYTMAIAARINALGVTQDVLMDNQKQIANKFNQALDQIQEGFKATHSAIATMQNAINQNAIALNQLVSELSNNFGSISATITEIVKRLDQLESDVQIDRLINGRMTALNTYVTQTLLDIKR

1 201 - 1 350
N107_S12  VEDQARLAIEIINECVKSQSLRRGFCGVDKHVYSILQRAPSGFMFVHVGIEFTDFVNITVAAGLCVNTERGEIGLLPKQGVFTPCYQDSDDGPWCVTPVEMLQLQNLTVENVIVGSCQKNFTYTNNSIDVNGNPTFNQSFVDELLEHYKK
N107_S15  VEDQARLAIEIINECVKSQSLRRGFCGVDKHVYSILQRAPSGFMFVHVGIEFTDFVNITVAAGLCVNTERGEIGLLPKQGVFTPCYQDSDDGPWCVTPVEMLQLQNLTVENVIVGSCQKNFTYTNNSIDVNGNPTFNQSFVDELLEHYKN

1 351 - 1 457
N107_S12  FTTLIPQLGISNITASLADVSQELGVLQEVVKQLNESYIKLEDLGKYEQYIKWPWYVWLGFIAGLVAIFLVLVMCLCATGCCTSCRGMCPKFRPRYEEFDYERLHTN
N107_S15  VTTLIPQLGISNITASLADVSQELGVLQEVVKQLNESYIKLEELGKYEQYIKWPWYVWLGFIAGLVAIFLVLVMCLCATGCCTSCRGMCPKFRPRYEEFDYERLHTN
```
