## Supplementary Table 2 for "*Ambecovirus*, a novel *Betacoronavirus* subgenus circulating in neotropical bats sheds new light on bat-borne coronaviruses evolution"

**Supplementary Table 1** **-** Descriptive statistics of complete and draft coronavirus genomes recovered in this study and metadata associated with the samples.

| **Pool name** | **Species** | **Nº specimens** | **Contig length** | **Total Pass Filter Reads** | **Mapped Read Count** | **Mean Coverage Depth** | **Sex** | **Age** | **Reproductive stage*** |
| --- | --- | --- | --- | --- | --- | --- | --- | --- | --- |
| N107_10 | *P. gymnonotus* | 1 | 5,334 | 2908050 | 5131 | 17.54 | Male | Adult | Active |
| N107_11 | *P. gymnonotus* | 1 | 10,882 | 3206766 | 13073 | 45.04 | Male | Adult | Inactive |
| N107_12 | *P. gymnonotus* | 1 | 30,422 | 3400908 | 101845 | 336.54 | Male | Adult | Inactive |
| N107_13 | *P. gymnonotus* | 1 | 8,477 | 1178662 | 1865 | 6.25 | Male | Adult | Inactive |
| N107_15 | *P. gymnonotus* | 1 | 30,414 | 3294633 | 182751 | 631.17 | Male | Adult | Inactive |
| N107_17 | *P. gymnonotus* | 1 | 4,139 | 3647063 | 1927 | 6.65 | Male | Adult | Inactive |
| N107_31 | *P. gymnonotus* | 1 | 23,261 | 3977163 | 37408 | 127.28 | Male | Adult | Active |
| N107_32 | *P. gymnonotus* | 1 | 12,301 | 3789009 | 21238 | 64.27 | Male | Adult | Active |
| N107_34 | *C. perspicillata* | 1 | 3,682 | 4793598 | 2147 | 7.32 | Female | Adult | Inactive |
| N152_004 | *P. gymnonotus* | 2 | 9,276 | 2428400 | 858 | 3.54 | Female/Male | Adult/Adult | Inactive/Active |
| N152_005 | *P. gymnonotus* | 2 | 11,358 | 3199622 | 1909 | 7.14 | Male | Adult | Active |
| N152_006 | *P. gymnonotus* | 2 | 2,032 | 3287243 | 306 | 1.29 | Male | Adult | Active |
| N152_007 | *P. gymnonotus* | 2 | 2,049 | 3496616 | 135 | 0.50 | Male | Adult | Active |
| N152_013 | *P. gymnonotus* | 2 | 11,366 | 3140953 | 2977 | 11.06 | Female/Male | Adult/Adult | Inactive/Inactive |
| N152_014 | *P. gymnonotus* | 2 | 15,974 | 755285 | 1262 | 5.24 | Male/Male | Adult/Adult | Inactive/Active |
| N152_016 | *P. gymnonotus* | 2 | 14,587 | 3216751 | 8191 | 30.13 | Male/Male | Adult/Subadult | Active/Inactive |
| N152_018 | *P. gymnonotus* | 2 | 2,112 | 1802108 | 272 | 0.90 | Male/Male | Adult/Adult | Active/Inactive |
| N152_080 | *P. gymnonotus* | 1 | 1,207 | 1002122 | 65 | 0.20 | Male | Adult | Active |
| N152_160 | *P. gymnonotus* | 1 | 1,705 | 1152208 | 103 | 0.38 | Female | Adult | Lactating |

* Female individuals were classified into three categories: pregnant, lactating, or inactive (non-reproductive). Males were considered reproductively active when they exhibited visibly enlarged testes.
